# Food webs can deliver win-win strategies for tropical agroforestry and biodiversity conservation

**DOI:** 10.1101/2024.05.31.596784

**Authors:** Crinan Jarrett, Luke L. Powell, Tabe Tiku Regine Claire, Cyril Kowo, Diogo F. Ferreira, Alma L.S. Quiñones, Andreanna J. Welch, Daniel T. Haydon, Jason Matthiopoulos

## Abstract

Balancing biodiversity conservation and agricultural productivity is commonly regarded as a trade-off, but such analyses overlook ecosystem services that functional biodiverse communities provide in agroecosystems, and the possibility that win-win strategies may exist. We developed a dynamic mechanistic community model of the bird-insect food web associated with African cocoa agroforestry, structurally informed by metabarcoding data on bird diets, and fitted to trapping data on species abundances. We used the model to predict equilibrium community composition under varying intensities of shade management and pesticide use. Our results indicate that low-intensity farming favours forest bird species, and potential pollinator abundance, with no increase in pest biomass. Furthermore, using simulations of pesticide application, we found that pesticides do not effectively reduce pest biomass, and result in forest bird extinction. Our mechanistic framework combines the influence of management and the direct and indirect effects of species’ interactions, and demonstrates that low intensity agriculture may provide a win-win for biodiversity and ecosystem services.

## Introduction

Low intensity small-holder agriculture is widely practiced in the tropics, where some of the world’s economically-poorest but biodiversity-richest regions are located (Tscharntke et al., 2012). In these areas, farmers commonly live on <1€/day, surrounded by some of the highest levels of wildlife diversity on Earth (Tscharntke et al., 2012). With limited resources to manage their land, farmers in tropical regions often rely heavily on hand-held tools and ecosystem services provided by wildlife (Tscharntke et al., 2011a). Therefore, low-intensity agriculture offers opportunities for both agricultural productivity and biodiversity conservation (Clough et al., 2011; Tscharntke et al., 2011b). However, it remains unclear how to manage these systems to achieve optimal combinations of the ostensibly competing aims of sustainable agricultural yields and biodiversity conservation.

The growing demand for food is pressurising policymakers into encouraging agricultural expansion and intensification (Ordway et al., 2017). Consequently, effectively managed wildlife-friendly agriculture is more important now than ever (Tscharntke et al., 2012). Intensification of agriculture commonly results in the replacement of floristically diverse agroecosystems for monocultures, often treated with high levels of chemical inputs (Clough et al., 2009; Tscharntke et al., 2011b). This intensification results in community changes, loss of biodiversity and abundance declines (Cassano et al., 2009; De Beenhouwer et al., 2013; Ferreira, Darling, et al., 2023; Jarrett et al., 2021). Changes in wildlife communities could also lead to unexpected loss of agricultural productivity, for instance due to reductions in animal groups that provide ecosystem services, such as pollination and pest control (Maas et al., 2013, 2016).

Understanding the trade-offs and synergies between biodiversity and productivity is complex, and has rarely been approached from a community-wide perspective (but see Karp & Daily, 2014; Kean et al., 2003; Kross et al., 2011). Animal communities in low-intensity agroecosystems are diverse and contain species that influence productivity and biodiversity conservation outputs in different ways, both direct and indirect (Bagny et al., 2018; Maas et al., 2013; Toledo-Hernández et al., 2021). Species that affect productivity either directly (e.g., pollinators) or indirectly (e.g., predators of pollinators or pests) are broadly considered to provide ecosystem services or dis-services. Different community compositions therefore result in different outcomes for biodiversity conservation, ecosystem services and dis-services in agroecosystems. Ideally, systems would be managed to optimise a combination of desirable outcomes, for instance high abundance of species of conservation value and ecosystem service providers, and low abundance of dis-service providers. However, because each species can respond differently to habitat management, creating a potential cascade of direct and indirect effects of management on the whole food web, responses of communities to agricultural management are hard to predict.

Previous research into the effect of agricultural management on biodiversity or productivity has mostly focussed on establishing correlations between the observed abundance of different groups and management covariates (Blaser et al., 2018; Clough et al., 2011). The limitation with these correlative analyses is that they commonly assume that species respond to management independently from each other. In other words, a correlation is established between the observed density of a species and management covariates, without accounting for the fact that the densities of other species present may be at least as important in determining community states (Gotelli & Ellison, 2006; Janssen & Rijn, 2021; Kawatsu et al., 2021; Tylianakis et al., 2007). To more accurately predict the effects of management on wildlife populations, and consequently on biodiversity conservation, ecosystem service and dis-services, we need a framework that can incorporate both direct responses of species to management and interactions between species in the food web.

Mechanistic community models could provide this framework. However, the common issue with such models is that they involve many parameters, and are consequently hard to fit to short-term field study data (Burt et al., 2018; Curtsdotter et al., 2019; Kawatsu et al., 2021). Community models are typically believed to require long time-series of data on ecological communities in order to distinguish the noise (demographic and environmental stochasticity) from the parameters governing the composition of communities (Ellner et al., 2002; Yodzis, 1998). However, dynamical models (e.g., both difference and differential equation models) focus on rates of change, not absolute abundances. Information on rates of change does not necessarily require long time-series: pairs of successive observations may be sufficient (Ives et al., 2003). Here, we sought to investigate fitting complex mechanistic community models to short time-series by exploiting a space-for-time substitution; we considered paired observations from a range of sites with shared parameters, so that community composition at time *t* depended only on community composition at time *t*-1. This experimental design is common in ecological studies, making our modelling framework applicable to datasets derived from typical 2-3 year long field projects, and particularly suitable for pooling multiple data fragments into a common inferential platform.

We modelled complex community dynamics using a set of coupled difference equations, based on concepts from traditional community ecology, but formulated as a series of GLMs, enabling us to directly fit them to field data using Bayesian methods. Our model incorporates data on species’ abundances and considers rates of change in the density of each species as a function of environmental covariates (including management). We informed trophic links between species through a dataset of >300 diet samples from insectivorous birds, processed using diet metabarcoding. This is, to our knowledge, the largest high-resolution dataset on Afrotropical bird diets, and provides novel and robust data on trophic connections between birds and arthropods in our system. Our models capture full-system dynamics, incorporating both the direct and indirect effects of management and species on each other.

Agroforestry, where crops are grown under a canopy of shade trees, provides an ideal model system to investigate community dynamics, and resulting ecosystem services and disservice outcomes. Such systems are a common form of agricultural production in tropical regions, can be highly diverse, and contain many service and disservice-providing species. Within agroforestry, the insectivore-insect food web is central and reflects important trade-offs between the conservation of vulnerable taxa, and the provisioning of ecosystem services and dis-services; insectivores, especially avian insectivores, are common in agroforestry (Jarrett et al., 2021, 2022) and can play an important role in pest control (Ferreira, Jarrett, et al., 2023; Maas et al., 2016). Additionally, insectivores are vulnerable to habitat degradation and consequently should be a priority for conservation in these landscapes (Bregman et al., 2014; Jarrett et al., 2021; Powell et al., 2015). Insects are a widespread group in agroforestry, and can be beneficial (pollination, pest control); however, they can also be extremely damaging to crops, causing crop losses of up to 40% (Akesse-Ransford et al., 2021; Bisseleua et al., 2013; Wessel & Quist-Wessel, 2015).

Here, we aimed to a) estimate long-term food web composition (community states) by capturing the mechanisms driving it, b) investigate the effects of management on densities of taxonomic groups in the food web, and assess the consequences for biodiversity conservation and ecosystem (dis-)services and c) predict changes in food web composition under simulated pesticide application scenarios.

## Methods

### Conceptual background

We assumed that communities of species fluctuate around a given (potentially unobserved) equilibrium state (Fig. 1). We use the term ‘species’ to describe taxonomic groups, which may be species but could also be functional groups, families, etc. We define community state as the vector of the abundances of all species at a given point in time 𝑵_𝑡_ = {𝑁_𝑡1_, … , 𝑁_𝑡𝑖_, … , 𝑁_𝑡𝐼_ }, where *I* is the total number of species, including those that happen to have a zero density in any particular system. While stable equilibria cannot be assumed to be observed in the data (Fig. 1), they are latent states towards which a system will tend, so they are the objective of our estimation.

**Figure 1.**
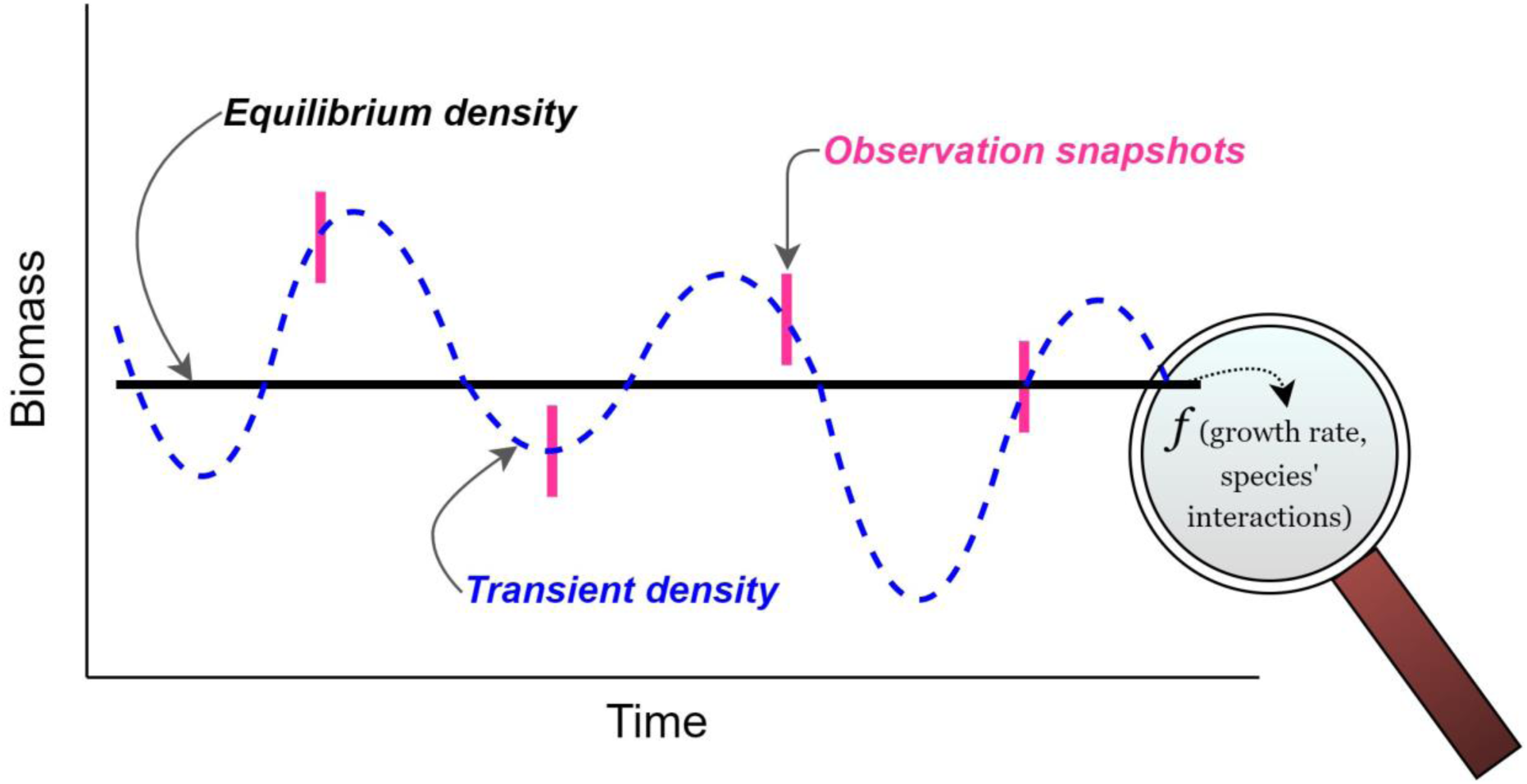
Rationale of our community model: we assume that the biomass of each species fluctuates stochastically (dashed blue line) around an equilibrium (solid black line), and this equilibrium density is determined by species’ intrinsic growth rates (which are a function of environmental covariates) and inter- and intra-specific interactions. When we observe species’ abundances, we see a snapshot of the transient densities (dashed blue line), but observation is inevitably made with error (represented by pink strips).

Community states are the result of species’ growth, environmental conditions and interactions between species. Observations of the state of the community were made at discrete time points, and captured a state of transience around the equilibrium (Fig. 1). Given sufficient observations of different transient states, we should be able to differentiate the noise (demographic and environmental stochasticity), from the parameters governing the equilibrium states. Our model was designed to fit a set of paired time-points (time *t* and *t*+1), and these pairs could come from different locations characterised by different environmental and management regimes, but were assumed to be governed by shared parameters. Therefore, species’ intrinsic growth rates and interactions were assumed to be constant across sites, whilst species’ densities could vary between sites.

### General model structure

Our model was based on a discrete-time Lotka-Volterra community model with *I* potentially interacting species. We modelled biomass of each species (a continuous variable) with a Gamma distribution where variance was equal to the mean (Eq. 1). The mean was written as a function of biomass at the previous time-step and the exponential of a linear predictor 𝐿_𝑡𝑖_, representing per-capita rate of change for each species at a given time.

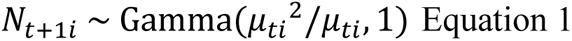

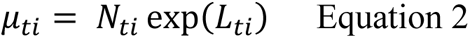

The log of the per-capita rate of change 𝐿_𝑡𝑖_ was written as a linear predictor, with an intercept 𝑏_𝑖_ interpreted as the log of the species-specific intrinsic growth rate (i.e., rate of change in the absence of all other species, whether predators or prey), and the sum of the effects of all interactions between species.

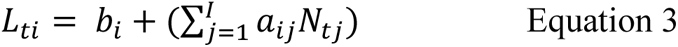

The coefficients 𝑎_𝑖𝑗_quantify the effect of the abundance of species *j* on the per-capita population growth rate of species *i.* Intraspecific effects (e.g., density dependence) were captured by 𝑎_𝑖𝑖_. We assumed a Holling Type I functional response between predators and prey to simplify model parametrisation, but non-linear functional responses could in theory be used here via quadratic terms or other common extensions of linear predictors.

In these coupled equations, the equilibrium state was given when exp(𝐿_𝑡𝑖_) = 1 for all *i*, and therefore 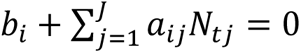 for all *i*. The equilibrium state can be found by multiplying the inverse of the 𝐀 matrix (containing the 𝑎_𝑖𝑗_ elements) by the vector of intrinsic growth rates 𝐛 (containing the 𝑏_𝑖_ elements).

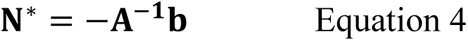

The elements (−𝑎_𝑖𝑗_^−1^) of the inverse of the **A** matrix (−𝐀^−𝟏^) capture the overall effect of each taxon on each other, defined as the change in equilibrium biomass of one taxon as a function of sustained increase in the biomass of another (Yodzis, 1988). These parameters therefore represent the combined influence of all direct and indirect effects of changes in biomass of species *j* or species *i*.

### Environmental covariates

We assumed that the log of the intrinsic growth rate (𝑏_𝑖_) of species *i* could be influenced by environmental covariates.

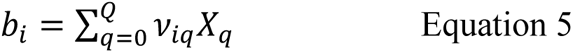

The linear predictor comprised 𝑄 covariates, 𝑋_𝑞_, affecting growth rate, and their respective regression coefficients 𝜈_𝑖𝑞_, where *q* refers to the *q*th covariate (the intercept 𝜈_𝑖0_was included by setting 𝑋_0_ = 1). Including the effect of environmental covariates on 𝑏_𝑖_ resulted in a **b** vector whose elements were functions of these covariates. Consequently, equilibrium densities as calculated by Equation 4 changed linearly with environmental covariates.

Before fitting the community model to field data, we tested it on simulated data to assess the accuracy and precision of parameter posteriors (Appendix I).

## Application to cocoa agroforestry

### Data collection

We applied our model to field data collected from 28 Cameroonian cocoa farms during Jan-Feb and Aug-Sept 2019-2020. The farms were on a gradient of shade cover, ranging from 19% to 99%. Each farm was visited 2-4 times, and on each occasion, we surveyed birds and arthropods, and collected bird faecal samples for diet analysis. We aggregated species into groups, to limit the number of parameters to be estimated. Based on our field data, we determined the main groups that formed the bird-arthropod food web in these cocoa farms (Fig. 2). For birds, 60% of all insectivorous individuals captured belonged to one of 5 genera or belonged to a guild of forest specialists (Jarrett et al., 2021), so we clustered our food web accordingly (Appendix II). The genera *Camaroptera*, *Hylia*, *Platysteira* (wattle-eyes) and *Terpsiphone* (flycatchers) are small passerine birds, some species of which are sensitive to habitat degradation (Jarrett et al., 2021; Appendix II). *Ispidina* is a genus of small insectivorous kingfishers, considered habitat generalists (Jarrett et al., 2021; Naidoo, 2004). We classified arthropods as either ‘pests’ or ‘non-pests’ and then grouped them by order, except for Brown capsid (*Sahlbergella singularis*), the primary pest of cocoa in Africa (Bagny et al., 2018), which was included at species level.

**Figure 2.**
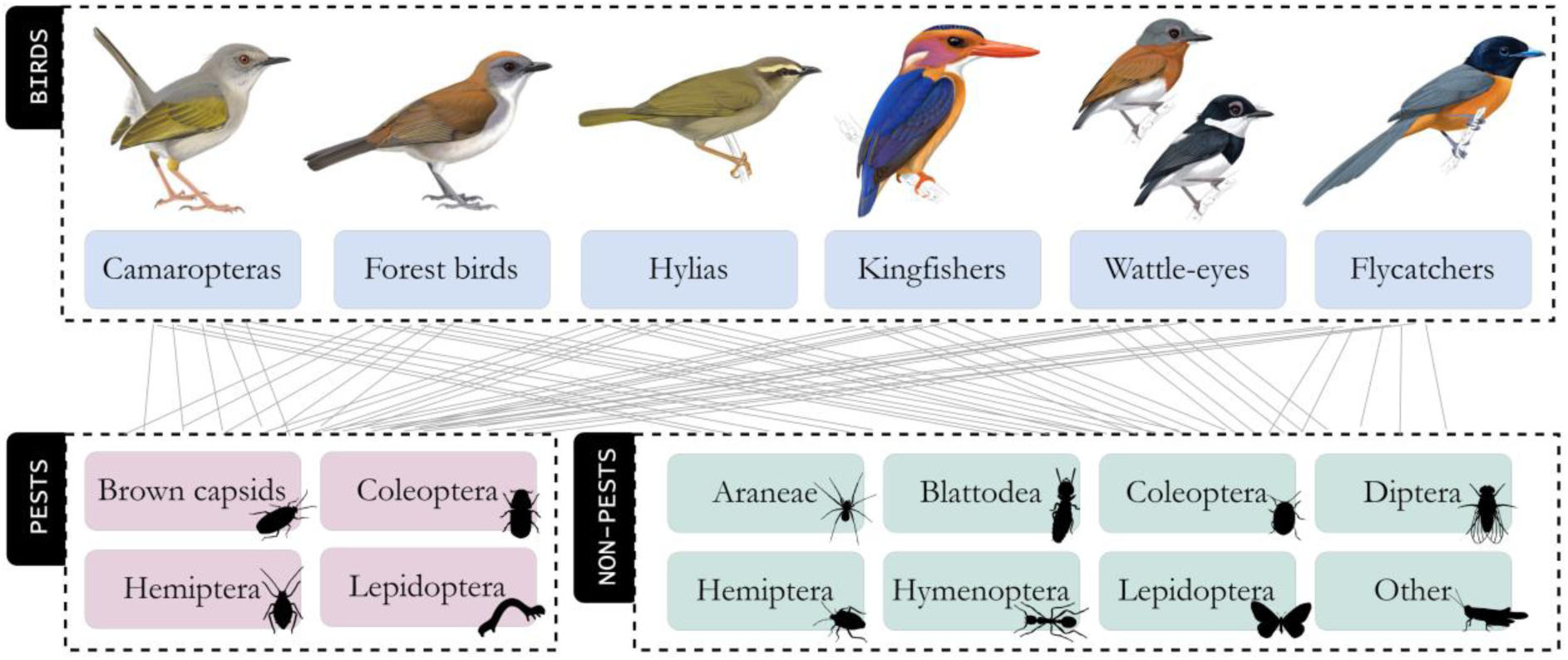
Structure of cocoa farm bird-arthropod food web: two trophic levels representing insectivorous birds (predators) and arthropods (prey). Bird illustrations by Faansie Peacock.

<u>Birds:</u> The bird dataset used for the model was that of Jarrett et al. (2022). The dataset consists of bird mist-net captures and simultaneously collected acoustic recordings from Cameroonian cocoa farms from Jan-Feb and Aug-Sept 2019 – 2020 (a period of 19 months). All birds caught were identified to species level and weighed, to allow conversion to biomass.

<u>Arthropods:</u> The arthropod dataset used for the model was identical to that Jarrett et al. (2023). It included count data for arthropods collected using three common survey methods: sweep-netting, malaise traps and visual surveys. Arthropods were identified to order level, except for brown capsid (*Sahlbergella singularis*), the primary pest of cocoa in Africa which was identified to species level. Aside from brown capsid, we considered three other potential pest groups, Coleoptera, Hemiptera and Lepidoptera – the distinction between pests and non-pests in these orders was made in the field based on empirical observations of individuals damaging cocoa crops (Jarrett et al., 2023).

<u>Trophic links:</u> we analysed faecal samples of 311 insectivorous birds (n=74 camaroptera, n=56 flycatcher, n=28 wattle-eye, n=65 kingfisher, n=29 hylia, n=59 forest birds) using diet metabarcoding methods to inform trophic links in the food web model (Appendix III).

<u>Covariate data:</u> the 28 cocoa farms sampled were distributed across gradient of shade cover. Our method for quantifying shade cover (in short, as percentage of the sky obscured by shade trees above the cocoa) is described in Jarrett et al. (2021).

### Observation models

To account for imperfect detection in field data, we used the observation models in Jarrett et al. (2022, 2023) for birds and arthropods respectively (for details see Appendix IV). The bird model combined mark-recapture data from mist-net captures with detections from acoustic recorder into a joint likelihood, to estimate population size at each site. In this model, site is considered to be the 1.5 Ha area covered by the sampling radius of the mist-nets, and thus the estimated bird population represents the number of birds whose home ranges overlap with this area (Jarrett et al., 2022). We then extrapolated bird population size to 1 km^2^.The insect model also integrates data from different sampling methods, in this case malaise traps, sweep-nets and visual surveys, assuming that each method sampled a population that was proportional (but not necessarily equal) to the true underlying population size (Jarrett et al., 2023; Miller et al., 2019). The model incorporated different capture rates for each method and taxon, to allow for varying detection. The estimated population size from this model was at the scale of one tree; we then extrapolated to 1 km^2^ (to match bird spatial scale) by assuming one tree occupies 9 m^2^ (Appendix IV).

We calculated the mean and standard deviation of the MCMC chains for each model state (bird or arthropod population size at farm *f = 1…28* and timepoint *t = 1…19*), and converted these to biomass by standardising by body mass of taxa (Appendix IV). We provided both the mean and standard deviation as data to the process model, thus propagating the uncertainty from the observation models.

### Process model parametrisation

We modelled community dynamics of birds and arthropods in cocoa as described in equations 1 – 5. The number of interacting taxa (*I*) was 18 (Fig. 2).

We modelled the log of the intrinsic growth rate of each taxon *i* as a function of shade cover at each farm *f* and the season during which timepoint *t* fell (Eq. 6; categorical variable:

𝑠𝑒𝑎𝑠𝑜𝑛_𝑡_ = 0 for dry season and 𝑠𝑒𝑎𝑠𝑜𝑛_𝑡_ = 1 for wet season). This assumes that a species caught in two different farms with equal shade cover and during the same season would have equal intrinsic growth rates. We constrained 𝑏_𝑖𝑓𝑡_ to be positive for arthropods to replicate the common food web parametrisation for primary producers, in the absence of lower trophic levels (Table 1; Pimm, 1982) and negative for birds (by specifying -1 * Gamma distribution; Table 1). Parameters describing the effects of covariates on the log of intrinsic growth rates were given normally distributed priors centred around 0 (Table 1), allowing for the effect of shade and season to be positive or negative.

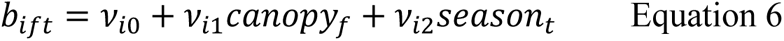

**Table 1.**
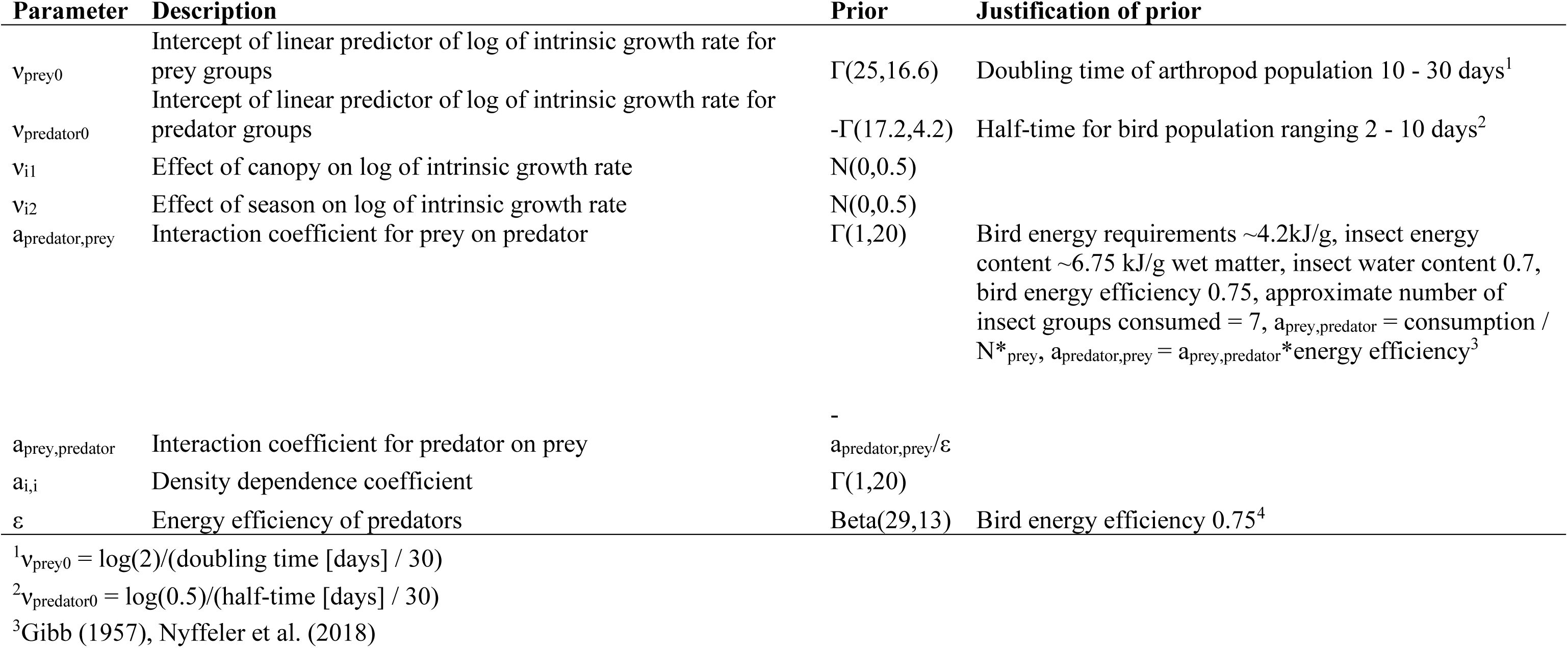
Description of parameters estimated by field data model, including model priors and the justification for prior distribution.

For the inter-specific interaction parameters, we assumed no direct competition, which seemed reasonable given differences in foraging niches among species (Appendix II). Whilst we included no direct interactions within trophic levels, this does not exclude indirect effects, which are captured by the model in the inverse of the interaction matrix (Eq. 4).

Of the 72 potential predator-prey interaction parameters (12 prey x 6 predators), we set 25 to 0 based on evidence from diet metabarcoding data from birds, that indicated minimal trophic links between certain bird genera and insect orders (Appendix III). Importantly, for all other interactions we did not consider information from diet metabarcoding, because quantifying interaction strength based on such diet data is complicated due to primer bias and other nuances of metabarcoding methods (Deagle et al., 2019). Instead, we modelled the remaining consumption parameters 𝑎_𝑝𝑟𝑒𝑑𝑎𝑡𝑜𝑟,𝑝𝑟𝑒𝑦_ using Gamma priors, assuming the effect of prey on predator growth rate was positive (Table 1). We restricted 𝑎_𝑝𝑟𝑒𝑦,𝑝𝑟𝑒𝑑𝑎𝑡𝑜𝑟_ to negative values and scaled with an energy efficiency coefficient (Table 1), to represent the negative effect of predators on the growth rate of prey, and the process of energy exchange from prey to predator.

Modelling community dynamics in discrete time can be complicated by the lack of instantaneous feedback loops and instabilities caused by the implicit time-lags in the discrete time formulation (Caswell & Neubert, 2006). This can cause explosive model behaviour and consequently, difficulty with fitting the model to data. Whilst our data were collected approximately every six months (Jan-Feb and Aug-Sept), we fit the model to linearly interpolated monthly time-steps to avoid such difficulties. This resulted in 19 total time-steps, spanning January 2019 to August 2020.

We ran the process model with 3 chains for 50,000 MCMC iterations (plus 10,000 burn-in). We fit all models using Bayesian inference with the JAGS 4.3.0 software (Plummer, 2017) executed using the runjags package (Denwood, 2016) in the R statistical computing environment (R Core Team, 2022).

### Community composition projections

To understand long-term effects of shade cover on community composition of birds and arthropods, we used the posteriors of parameters for growth rate and species interactions. We drew 1,000 MCMC equally spaced trials from our total set of MCMC iterations approximating the joint posterior, and used these draws to simulate community trajectories under different shade cover values over a 20 year period, following equations 1-3 and 6. For starting point, we took the mean value of density for each taxon estimated from the observation models.

### Arthropod suppression

We wished to investigate the long-term community-wide effects of artificially suppressing arthropod populations, as might occur through the application of pesticides or other control measures. Using the parameters estimated from fitting the community model to data, we simulated trajectories over a 20-year period, as above, but under different scenarios of arthropod population suppression: we simulated 9 scenarios, ranging from 10% to 90% impacts on arthropod intrinsic growth rates. This broad range of potential pesticide effects was chosen because pesticide-induced mortality can vary according to the product and the taxon (e.g., from empirical studies, 0.07 to 0.98 survival rate after 1-day chemical application, Janssen & Rijn 2021). We compared the arthropod suppression scenarios with a scenario with no perturbation, for which we used the posteriors from our field data; therefore, we assume that chemical application in our field sites was negligible, based on information provided by the farmers. We assumed that chemical application, and therefore growth rate reduction, occurred once per year, and the other timesteps in the year were parametrised using the original parameter posteriors with no reduction. Additionally, we assumed that all arthropods were affected equally by pesticide application; this is likely a simplification of what occurs in the field, but nonetheless provides useful insight into the food-web-wide effects of arthropod declines.

## Results

### Shade cover affects equilibrium food web composition: more forest birds and fewer brown capsids under dense shade

Over a 20-year period, communities settled at different compositions according to shade cover (Fig. 3): at low shade cover, the bird community was dominated by kingfishers, followed by camaroptera and forest birds. At high shade cover, bird communities were dominated by forest birds, with at least two-times more biomass in this group than any other bird group. Under all shade scenarios, wattle-eyes, hylia and flycatchers occurred at low biomass, though flycatchers and hylia showed a slight decrease in biomass with increasing shade, and wattle-eyes the opposite trend.

**Figure 3.**
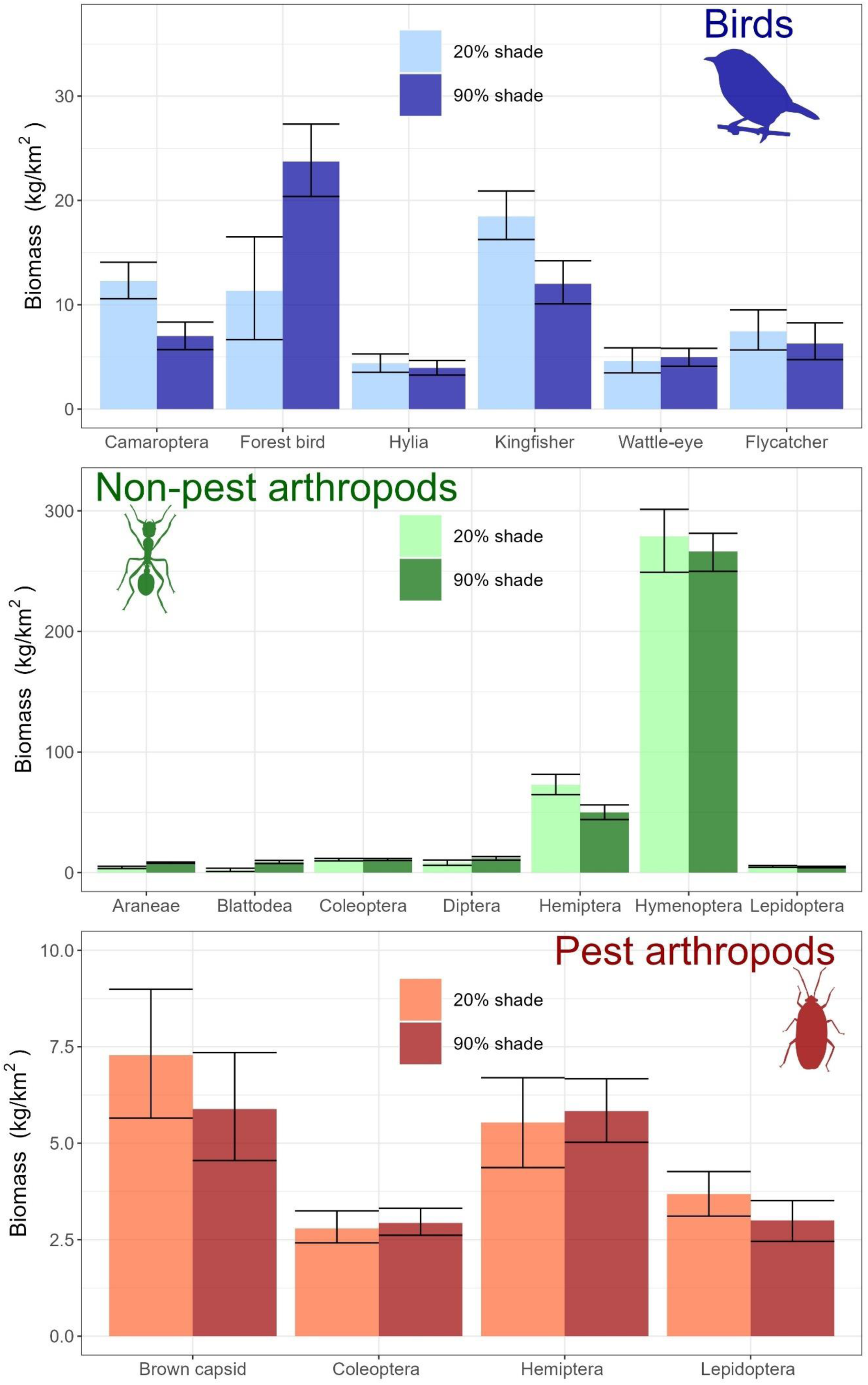
Biomass of taxa after 20 years under low (20%) and high (90%) shade cover. Biomass was calculated at each timepoint using Eq. 1-3 & 6, with parameters estimated from fitting the model to field data. The final timepoint, presented here, represents biomass during the dry season. Bars represent mean biomass values and error bars capture 95^th^ quantiles.

Non-pest arthropod communities under low shade had higher biomass of Hemiptera, Hymenoptera and Lepidoptera, whilst all the other groups showed higher biomass under high shade cover. Araneae and Blattodea showed the steepest increase in biomass with shade cover. For the pest community, under low shade biomass of capsids and lepidopteran pests was higher, and biomass of hemipteran and coleopteran pests was marginally lower.

After a 20-year period, biomass of brown capsids was on average 7.5 kg/km^2^ (0.08 kg/Ha) on sunny farms, and 6 kg/km^2^ (0.06 kg/Ha) on shady farms (Fig. 3). In contrast, forest bird biomass on sunny farms was on average 12 kg/km^2^ (0.12 kg/Ha) whilst on shady farms it was 24 kg/km^2^ (0.24 kg/Ha; Fig. 3).

### Changes in food web composition are mediated through interactions between taxa as well as shade management

The food web composition under different shade cover values was a result of both the intrinsic response of taxa to shade cover (for νi1 posterior summary see Appendix V) and interactions between taxa (Fig. 4; Appendix V). Net effects of each taxon on each other were captured in the inverse of the interaction matrix (-**A**^-1^): in general, the net effects of birds on arthropods were negative, and varied in strength. One of the strongest effects was between forest birds and Hymenoptera (Fig. 4). In some cases, there were positive interactions between birds and arthropods; these usually occurred between pairs where the direct trophic interaction had been set to 0 based on evidence from diet data, for instance between forest birds and Diptera. In these cases, as there is no predation, an increase in the biomass of the bird taxon arises through indirect interactions (for example by minimising competition for the non-prey taxon, Fig. 4). Whilst direct effects amongst bird taxa and amongst arthropod taxa were set to zero in the model, net effects between bird taxa were all negative, whilst between arthropod taxa there were some positive effects.

**Figure 4.**
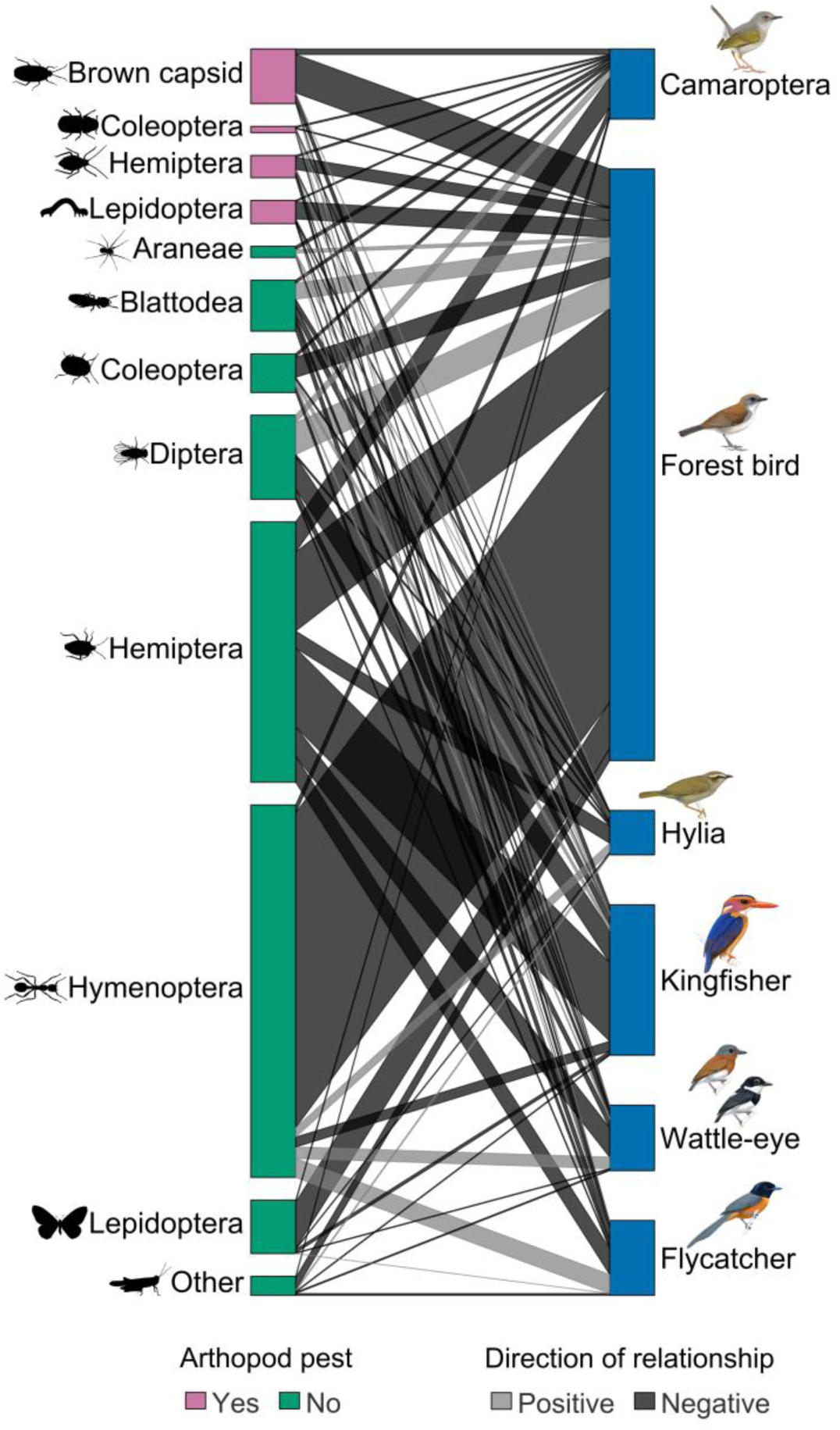
Interactions between bird and arthropod groups in the food web. Represented is the net effect of bird groups on arthropod groups, i.e., the change in equilibrium biomass of one taxon caused by the addition of biomass of another taxon, estimated by inverting the interaction matrix (-**A**^-1^). Bird illustrations by Faansie Peacock.

Importantly, in some cases, the difference in biomass between sunny and shade farms was more strongly influenced by interactions than by intrinsic responses to shade. For instance, the effect of shade cover on the growth rate of hylia was negative (Appendix V), but community states showed almost no change in biomass with shade, indicating that interactions with other taxa (e.g., availability of prey, reduced competition) counteracted the direct effect of shade on growth rate.

### Pesticide application is least effective at suppressing brown capsids in shady farms and results in forest bird extinction

The long-term effect of simulated pesticide application on biomass of taxa varied according to shade cover (Fig. 5). For brown capsids in low shade farms, a 60% pesticide intensity resulted in a decrease in biomass of -46% (equivalent to 3.3 kg/km^2^), whilst the equivalent application in high shade farms resulted in a decrease of -30% (2 kg/km^2^). For forest birds, the negative effect of pesticide application was exacerbated in low shade: in sunny farms forest birds became extinct at >40% pesticide intensity, whilst in shady farms they persisted up to 80% pesticide intensity. Other groups showed varying responses: for instance, camaroptera showed a steeper decrease in biomass with pesticide intensity under shady conditions, whilst other groups such as Araneae and hylia showed consistent declines in biomass with pesticide intensity independent of shade cover. Interestingly, blattodea was the only taxon that increased in biomass with pesticide application. Overall, to achieve a decline of less than 50% in non-pest taxa, pesticide application can only reach 10% in the sunniest farms, and 20% in shady farms (Fig. 5). This level of application achieves a maximal reduction in pests of 19% (Fig. 5).

**Figure 5.**
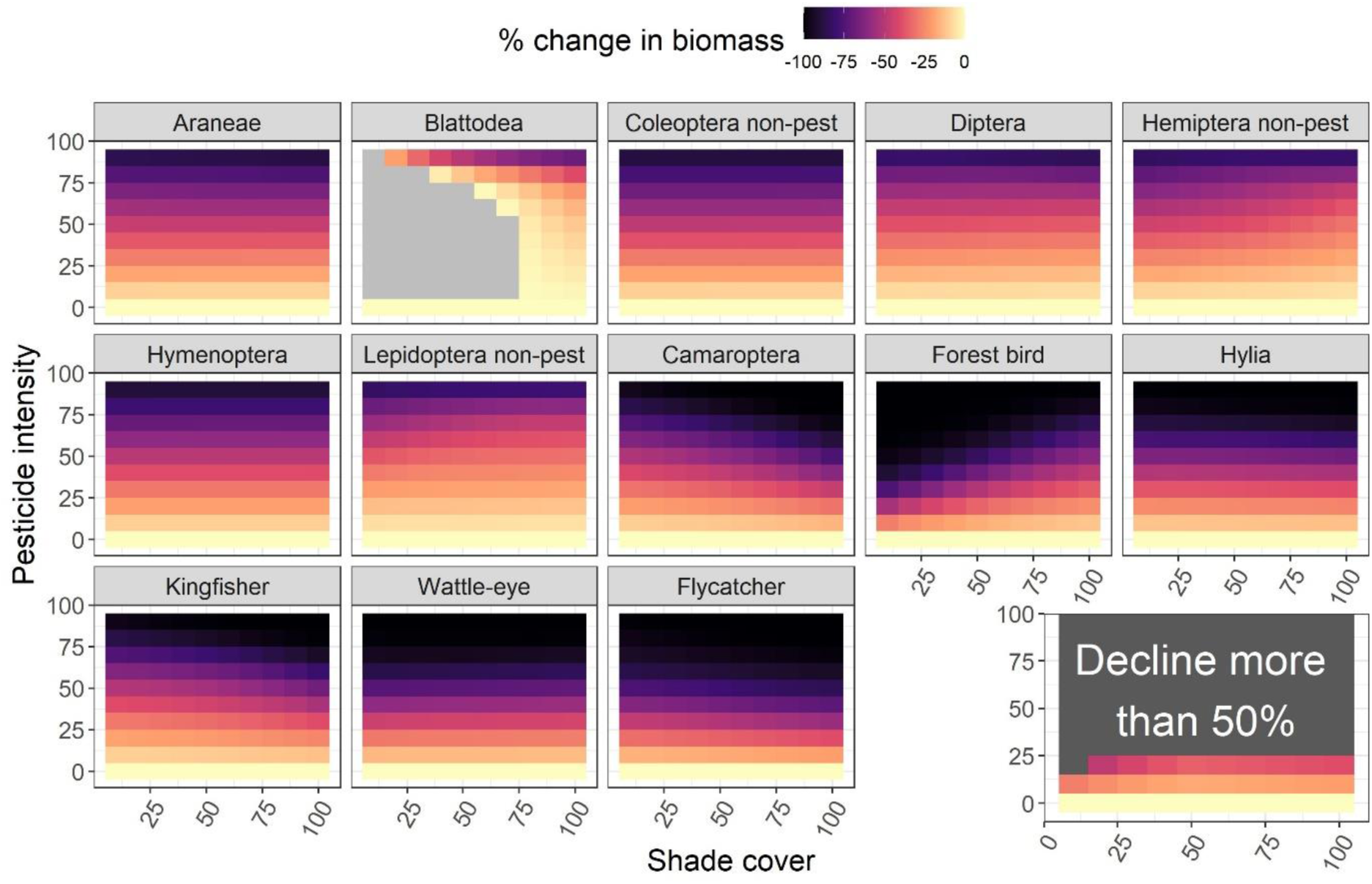
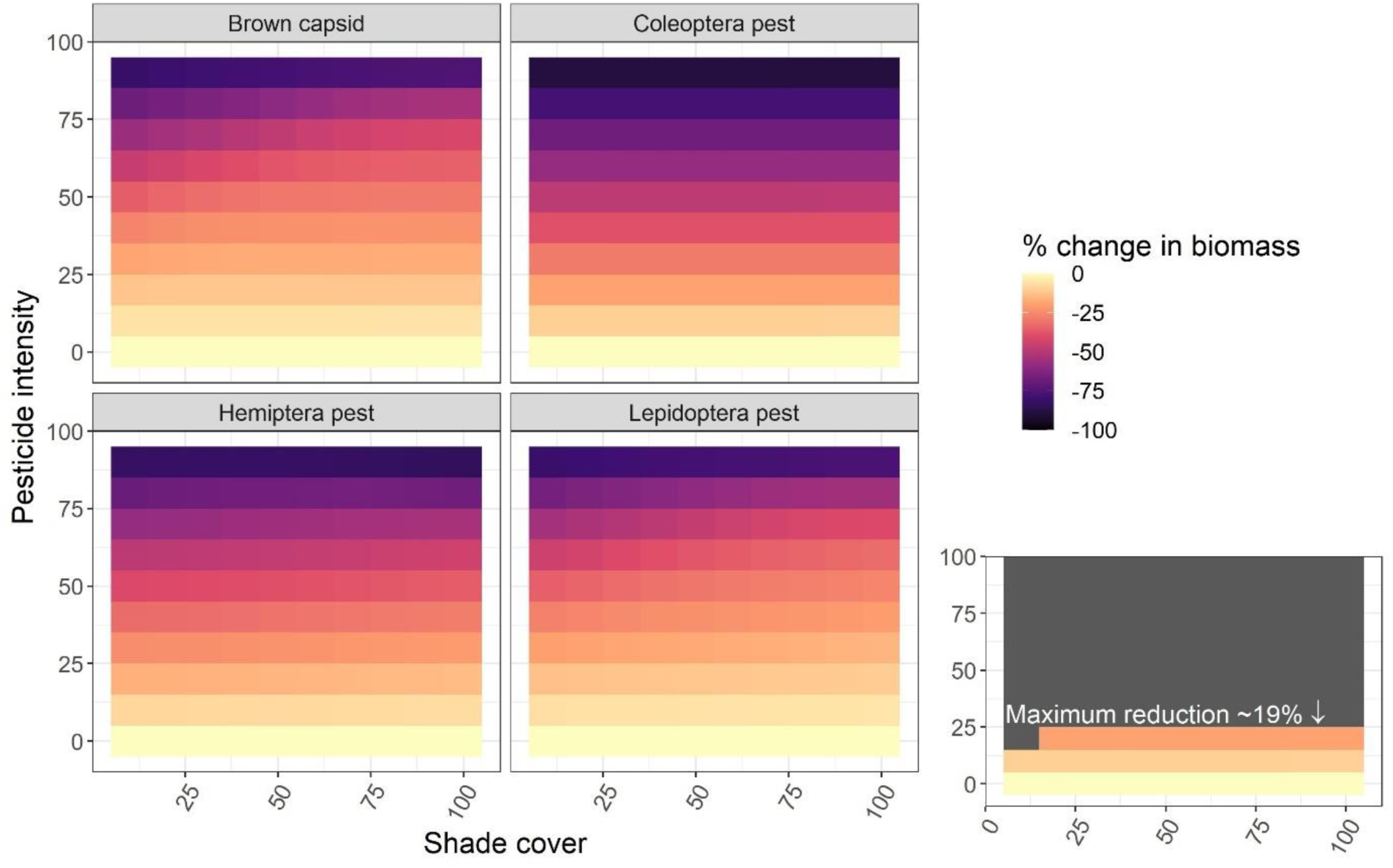
Reduction in biomass (%) in birds and non-pest arthropods (above) and pests (below) under a range of pesticide intensity and shade cover scenarios for each taxon. Lighter colours indicate less change, and darker ones more change. In one case there is positive change, which is indicated in light grey. The small inlaid panel above summarises the scenarios where the maximum decline amongst non-pest taxa is less than 50%. The inlaid panel below shows, for those same scenarios, the maximum reduction achieved in pest populations.

## Discussion

We investigated the long-term community-wide effects of management in agricultural habitats, using African cocoa agroforestry as a model system. We used a community modelling approach, which incorporated information on species’ interactions and responses to shade cover. Our results indicate important changes in community composition with management, revealing that low-intensity farming favours forest bird species and potential pollinators, with no increase in pest biomass.

Equilibrium community composition in sunny farms supports higher biomass of generalist bird taxa such as *Ispidina* kingfishers and camaropteras, and lower biomass forest birds. These findings are consistent with previous studies from the same study system, and other agroforestry systems around the world, indicating that cocoa farms, including intensively managed ones, can support relatively high diversity of birds, but lose specialised insectivores (Bennett et al., 2021; Faria et al., 2006; Jarrett et al., 2021; Waltert et al., 2005). However, here we go beyond commonly-used correlational approaches and predict states of communities in agroforestry from a mechanistic community model, and accounting for the influence of species interactions as well as the direct influence of shade cover. This reveals that the response of taxa to shade cover is driven both by an intrinsic response to shade (captured by the 𝜈_𝑖1_parameter) and the effect of other taxa in the community. For instance, hylia showed a likely negative effect of shade on growth rate, yet the population settled at similar biomass in shady farms compared to sunny ones, indicating a response to other taxa in the community. In contrast, forest birds show an intrinsic positive response to shade cover, reflecting perhaps habitat and microclimate requirements for breeding or foraging (Jirinec et al., 2022; Powell et al., 2015). Our findings suggest that widespread intensification and expansion of cocoa agriculture, as seen for instance in much of Côte d’Ivoire, will likely result in a landscape dominated by generalist bird taxa, and devoid of forest birds and other sensitive insectivores (Kupsch et al., 2019).

The non-pest arthropod community in both sunny and shady farms was dominated by Hymenoptera, with slightly higher biomass in sunny farms. Diptera occurred at higher biomass in shady farms, as did Araneae and Blattodea, with the latter two showing the most extreme decline down to close to zero biomass in sunny farms. In general, therefore, shady farms supported higher biomass of potential pollinators (Diptera), natural enemies (Araneae), and other ecosystem service providers as found by Ambele et al. (2023). Lepidoptera and Hemiptera decreased in biomass with increasing shade, mostly driven by an intrinsic effect of shade cover on growth rate, perhaps reflecting microhabitat requirements for breeding or development. Our results suggest that widespread intensification of cocoa agroforestry into monocultures results in broad changes in arthropod community composition, driven by a combination of intrinsic responses to shade cover and inter-specific interactions. Intensified farms support arthropod communities with fewer potential pollinators and natural enemies, and near extinction of certain taxa; such changes are a reflection of the global trends of insect declines caused by agricultural intensification (Lister & Garcia, 2018; Outhwaite et al., 2022). Recent studies have shown an additional interactive effect between climate warming and agricultural intensification, so that in intensified landscapes, insect declines due to climate warming are exacerbated (Outhwaite et al., 2022). Together, these findings indicate the high risks associated with agricultural intensification. Especially in small-holder agriculture, where farmers have limited resources and depend heavily on services provided by wild fauna, a decline in important arthropod groups could lead to unsustainable losses.

In the pest community, brown capsids and lepidopteran pests persisted at higher biomasses in sunny farms, whilst hemipteran and coleopteran pests showed the opposite trend. However, these differences were relatively small with overlapping quantiles, potentially due to limited data on pest populations (brown capsids are notoriously hard to detect in the field). Our results indicated that both brown capsid and lepidopteran pests had likely lower growth rates in shady farms; additionally, both groups were consumed by several bird taxa, including forest birds and wattle-eyes, that were more abundant in shady farms. Biological control by native predators in our study sites likely constitutes an important service for farmers with limited resources for agricultural inputs (Ferreira et al., 2023). Previous studies have found higher density of brown capsids in sunny farms (Babin et al., 2010; Bagny Beilhe et al., 2018), and such trends could explain long-term yield declines seen in intensified African cocoa farms (Ahenkorah et al., 1987).

Under simulated pesticide application scenarios, assumed to reduce arthropod growth rates, we found that shade had an influence on the outcomes of these interventions. Medium levels of pesticide application in sunny farms resulted in the extinction of forest birds, likely due to reduction in prey populations; indeed, forest birds showed the highest susceptibility to chemical use amongst all taxa, becoming extinct even in shady farms at higher intensity applications. Arthropods themselves rarely became extinct except at the highest level of chemical application, but also responded varyingly according to shade. Importantly, applying pesticides in shady farms did little to suppress brown capsid populations, likely due to a die-off of predators permitting population persistence. Overall, this investigation suggested that pesticide use, especially in shady farms, is largely ineffective because it leads to the extinction of predators and consequent higher survival of arthropods. These findings suggest limitations of laboratory testing of pesticides; whilst in laboratory conditions pest populations may be easily eradicated by chemicals, outcomes may change in the presence of other taxa. Our results support previous modelling work indicating the effectiveness of pesticide application is strongly mediated through interactions with other taxa (Janssen & Rijn, 2021).

The findings from our pesticide simulations match an experimental study conducted in the same study sites, showing that insectivorous birds and bats greatly enhance agricultural yields but only under high shade conditions (Ferreira et al., 2023). These analyses both suggest that natural pest control by birds and bats may be highly effective in shady farms, whereas productivity will decline (Ferreira et al., 2023) and pest communities persist in their absence. Overall, recent research globally clearly supports the importance of low intensity agriculture and agricultural landscapes in the maintenance of ecosystem services (Bommarco et al., 2013; da Silva et al., 2024; Dainese et al., 2019).

Together, our results provide robust evidence that intensification of agroforestry results in declines in ecosystem service-providing arthropods and the high risk of forest bird extinction. We also provide a novel framework for ecosystem-level management, an often-aspired to but rarely achieved practice (Brussard et al., 1998; Duru et al., 2015). Here, we tested a relatively simplified version of the food web model. Extensions to this model would be to include more types of species interactions (e.g., pollination, or predation between arthropod groups), as well as additional predators (e.g., insectivorous bats). Structural features such as non-linear functional responses, prey switching and Allee effects could be explored using common extensions of our linear predictors (Asseburg et al., 2006; Lindmark et al., 2019). Such non-linear extensions could result in more equilibria, though this remains to be tested.

An important extension to the model would be to add crops as a node in the food web; this would allow direct estimation of changes in crop biomass as a function of management and food web composition. Our intention was to test this model in a simplified scenario to minimise the number of parameters required to be estimated, and therefore we clustered most species into higher taxonomic groups. Additionally, we simplified food-web structure by excluding direct interactions within trophic levels. This clustering and simplification may have led to increased uncertainty in the estimation of certain parameters. For instance, the effect of shade cover within arthropod orders may be highly variable and therefore hard to represent with one single parameter. More exploration is needed to investigate how data sufficiency compares with level of taxonomic aggregation. The less data there are, the more aggregated a model may need to be and the more noise the results will contain. Whether our existing dataset would support a more complex food web remains to be tested.

Here, we presented a novel method for investigating population dynamics of complex communities. Applying our method to data from wildlife communities in agroforestry revealed that communities in low-intensity farms are important both for biodiversity conservation and for productivity. This win-win in low-intensity agroecosystems sheds light on the risks of pursuing intensified agriculture in an era of biodiversity crisis.

## Acknowledgements

We thank the dozens of cocoa farmers for allowing us to work within their farms and for the support provided. We thank M. Tchoumbou, M.N.F. Elikwo, C.A. Wandji and V. Talla for the help collecting data. We thank both the Congo Basin Institute and the International Institute of Tropical Agriculture (IITA), especially Emmanuel Meye, for logistic support. CJ was funded by a Caledonian PhD scholarship from the Royal Society of Edinburgh and managed through the Carnegie Foundation of Scotland. LLP was supported by the European Union’s Horizon 2020 research and innovation program under grant agreement No 854248. DFF was supported by a FCT fellowship (SFRH/BD/138310/2018).

## Statement of authorship

CJ, LLP, AJW, DTH and JM conceived the study, CJ led the writing of the manuscript, DTH and JM supported modelling work, CJ, LLP, TTRC, CK and DFF collected the data, ALSQ and AJW conducted and designed the lab methods.

## Data accessibility statement

code for this manuscript is archived at https://doi.org/10.6084/m9.figshare.24866001.v1. Data used for this manuscript are already published in public repositories.

## References

Ahenkorah, Y., Halm, B. J., Appiah, M. R., Akrofi, G. S., & Yirenkyi, J. E. K. (1987). Twenty Years’ Results from a Shade and Fertilizer Trial on Amazon Cocoa ( Theobroma cacao) in Ghana. Experimental Agriculture, 23, 31–39. 10.1017/S0014479700001101

Ambele, C. F., Bisseleua, H. D. B., Djuideu, C. T. L., & Akutse, K. S. (2023). Managing insect services and disservices in cocoa agroforestry systems. Agroforestry Systems, 0123456789. 10.1007/s10457-023-00839-x

Asseburg, C., Harwood, J., Matthiopoulos, J., Smout, S., & Camphuysen, C. J. (2006). The functional response of generalist predators and its implications for the monitoring of marine ecosystems. In I. L. Boyd & S. Wanless (Eds.), Top Predators in Marine Ecosystems (pp. 262–274). Cambridge University Press. 10.1017/CBO9780511541964.019

Babin, R., ten Hoopen, G. M., Cilas, C., Enjalric, F., Gendre, P., & Lumaret, J.-P. (2010). Impact of shade on the spatial distribution of Sahlbergella singularis in traditional cocoa agroforests. Agricultural and Forest Entomology, 12, 69–79. 10.1111/j.1461-9563.2009.00453.x

Bagny Beilhe, L., Babin, R., & ten Hoopen, M. (2018). Insect pests affecting cocoa. In P. Umaharan (Ed.), Achieving sustainable cultivation of cocoa: Genetics, breeding, cultivation and quality. Burleigh Dodds Science Publishing.

Bagny, L., Babin, R., & Ten Hoopen, G. M. (2018). Insect pests affecting cacao. In P. Umaharan (Ed.), Achieving sustainable cultivation of cocoa (pp. 303–326). Burleigh Dodds Science Publishing. 10.19103/AS.2017.0021.19

Bennett, R. E., Sillett, T. S., Rice, R. A., & Marra, P. P. (2021). Impact of cocoa agricultural intensification on bird diversity and community composition. Conservation Biology, 36(1), e13779. 10.1111/cobi.13779

Bommarco, R., Kleijn, D., & Potts, S. G. (2013). Ecological intensification: Harnessing ecosystem services for food security. Trends in Ecology and Evolution, 28(4), 230–238. 10.1016/j.tree.2012.10.012

Brussard, P. F., Reed, J. M., & Tracy, C. R. (1998). Ecosystem management: What is it really? Landscape and Urban Planning, 40(1–3), 9–20. 10.1016/S0169-2046(97)00094-7

Burt, J. M., Tinker, M. T., Okamoto, D. K., Demes, K. W., Holmes, K., & Salomon, A. K. (2018). Sudden collapse of a mesopredator reveals its complementary role in mediating rocky reef regime shifts. Proceedings of the Royal Society B, 285(20180553). 10.1098/rspb.2018.0553

Caswell, H., & Neubert, M. G. (2006). Reactivity and transient dynamics of discrete-time ecological systems. Journal of Difference Equations and Applications, 11(4–5), 295–310. 10.1080/10236190412331335382

Clough, Y., Barkmann, J., Juhrbandt, J., Kessler, M., Wanger, T. C., Anshary, A., Buchori, D., Cicuzza, D., Darras, K., Putra, D. D., Erasmi, S., Pitopang, R., Schmidt, C., Schulze, C. H., Seidel, D., Steffan-Dewenter, I., Stenchly, K., Vidal, S., Weist, M., … Tscharntke, T. (2011). Combining high biodiversity with high yields in tropical agroforests. Proceedings of the National Academy of Sciences, 108(20), 8311–8316. 10.1073/pnas.1016799108

Curtsdotter, A., Thomas Banks, | H, Banks, J. E., Jonsson, M., Jonsson, T., Laubmeier, A. N., Traugott, | Michael, & Bommarco, R. (2019). Ecosystem function in predator-prey food webs-confronting dynamic models with empirical data. J Anim Ecol, 88, 196–210. 10.1111/1365-2656.12892

da Silva, L. P., Mata, V. A., Lopes, P. B., Pinho, C. J., Chaves, C., Correia, E., Pinto, J., Heleno, R. H., Timoteo, S., & Beja, P. (2024). Dietary metabarcoding reveals the simplification of bird-pest interaction networks across a gradient of agricultural cover. *Molecular Ecology*, March, accepted. 10.1111/mec.17324

Dainese, M., Martin, E. A., Aizen, M. A., Albrecht, M., Bartomeus, I., Bommarco, R., Carvalheiro, L. G., Chaplin-kramer, R., Gagic, V., Garibaldi, L. A., Ghazoul, J., Grab, H., Jonsson, M., Karp, D. S., Letourneau, D. K., Marini, L., Poveda, K., Rader, R., Smith, H. G., … Tschumi, M. (2019). A global synthesis reveals biodiversity-mediated benefits for crop production. 1–13.

Deagle, B. E., Thomas, A. C., McInnes, J. C., Clarke, L. J., Vesterinen, E. J., Clare, E. L., Kartzinel, T. R., & Eveson, J. P. (2019). Counting with DNA in metabarcoding studies: How should we convert sequence reads to dietary data? Molecular Ecology, 28(2), 391–406. 10.1111/MEC.14734

Denwood, M. J. (2016). runjags an R package providing interface utilities, model templates, parallel computing methods and additional distributions for MCMC models in JAGS. Journal of Statistical Software, 71(9), 1–25. 10.18637/jss.v071.i09

Duru, M., Therond, O., Martin, G., Martin-Clouaire, R., Magne, M. A., Justes, E., Journet, E. P., Aubertot, J. N., Savary, S., Bergez, J. E., & Sarthou, J. P. (2015). How to implement biodiversity-based agriculture to enhance ecosystem services: a review. Agronomy for Sustainable Development, 35(4), 1259–1281. 10.1007/s13593-015-0306-1

Faria, D., Laps, R. R., Baumgarten, J., & Cetra, M. (2006). Bat and bird assemblages from forests and shade cacao plantations in two contrasting landscapes in the Atlantic Forest of southern Bahia, Brazil. Biodiversity and Conservation, 15(2), 587–612. 10.1007/s10531-005-2089-1

Ferreira, D. F., Jarrett, C., Wandji, A. C., Atagana, P. J., Rebelo, H., Maas, B., & Powell, L. L. (2023). Birds and Bats Enhance Yields in Afrotropical Cacao Agroforests Only Under High Shade. Agriculture, Ecosystems & Environment, 345, 108325.

Janssen, A., & Rijn, P. C. J. van. (2021). Pesticides do not significantly reduce arthropod pest densities in the presence of natural enemies. Ecology Letters, 24(9), 2010–2024. 10.1111/ELE.13819

Jarrett, C., Cyril, K., Haydon, D., Wandji, A., Ferreira, D., Welch, A., Powell, L., & Matthiopoulos, J. (2023). Fewer pests and more ecosystem service-providing arthropods in shady African cocoa farms: Insights from a data integration study. Journal of Applied Ecology, 61(2), 304–315. https://besjournals.onlinelibrary.wiley.com/doi/full/10.1111/1365-2664.14563

Jarrett, C., Haydon, D. T., Morales, J. M., Ferreira, D. F., Forzi, F. A., Welch, A. J., Powell, L. L., & Matthiopoulos, J. (2022). Integration of mark-recapture and acoustic detections for unbiased population estimation in animal communities. Ecology, 103(10), e3769.

Jarrett, C., Smith, T. B., Claire, T. T. R., Ferreira, D. F., Tchoumbou, M., Elikwo, M. N. F., Wolfe, J., Brzeski, K., Welch, A. J., Hanna, R., & Powell, L. L. (2021). Bird communities in African cocoa agroforestry are diverse but lack specialised insectivores. Journal of Applied Ecology, 58(6), 1237–1247. 10.1111/1365-2664.13864

Jirinec, V., Rodrigues, P. F., Amaral, B. R., & Stouffer, P. C. (2022). Light and thermal niches of ground-foraging Amazonian insectivorous birds. Ecology, 103(4), 1–13. 10.1002/ecy.3645

Karp, D. S., & Daily, G. C. (2014). Cascading effects of insectivorous birds and bats in tropical coffee plantations. Ecology, 95(4), 1065–1074. 10.1890/13-1012.1

Kawatsu, K., Ushio, M., van Veen, F. J. F., & Kondoh, M. (2021). Are networks of trophic interactions sufficient for understanding the dynamics of multi-trophic communities? Analysis of a tri-trophic insect food-web time-series. Ecology Letters, 24(3), 543–552. 10.1111/ele.13672

Kean, J., Wratten, S., Tylianakis, J., & Barlow, N. (2003). The population consequences of natural enemy enhancement, and implications for conservation biological control. Ecology Letters, 6(7), 604–612. 10.1046/J.1461-0248.2003.00468.X

Kross, S. M., Tylianakis, J. M., & Nelson, X. (2011). Reintroducing threatened falcons into vineyards reduces bird-damage to wine grapes. Conservation Biology, 26(1), 142–149. 10.1111/j.1523-1739.2011.01756.x

Kupsch, D., Vendras, E., Ocampo-Ariza, C., Batáry, P., Motombi, F. N., Bobo, K. S., & Waltert, M. (2019). High critical forest habitat thresholds of native bird communities in Afrotropical agroforestry landscapes. Biological Conservation, 230, 20–28. 10.1016/j.biocon.2018.12.001

Lindmark, M., Ohlberger, J., Huss, M., & Gårdmark, A. (2019). Size-based ecological interactions drive food web responses to climate warming. Ecology Letters, 22(5), 778– 786. 10.1111/ELE.13235

Lister, B. C., & Garcia, A. (2018). Climate-driven declines in arthropod abundance restructure a rainforest food web. PNAS, 115(44), E10397–E10406. 10.1073/pnas.1722477115

Maas, B., Clough, Y., & Tscharntke, T. (2013). Bats and birds increase crop yield in tropical agroforestry landscapes. Ecology Letters, 16(12), 1480–1487. 10.1111/ele.12194

Miller, D. A. W., Pacifici, K., Sanderlin, J. S., & Reich, B. J. (2019). The recent past and promising future for data integration methods to estimate species’ distributions. Methods in Ecology and Evolution, 10(1), 22–37. 10.1111/2041-210X.13110

Naidoo, R. (2004). Species richness and community composition of songbirds in a tropical forest-agricultural landscape. Animal Conservation, 7(1), 93–105. 10.1017/S1367943003001185

Outhwaite, C. L., McCann, P., & Newbold, T. (2022). Agriculture and climate change are reshaping insect biodiversity worldwide. Nature 2022, 1–6. 10.1038/s41586-022-04644-x

Pimm, S. L. (1982). Food webs. The University of Chicago Press.

Plummer, M. (2017). JAGS: A program for analysis of Bayesian graphical models using Gibbs sampling. (Version 4.3.0).

Powell, L. L., Cordeiro, N. J., & Stratford, J. A. (2015). Ecology and conservation of avian insectivores of the rainforest understory: A pantropical perspective. Biological Conservation, 188, 1–10. 10.1016/J.BIOCON.2015.03.025

R Core Team. (2022). *R: a language and environment for statistical computing* (Version 4.2.1). https://www.r-project.org/

Toledo-Hernández, M., Tscharntke, T., Tjoa, A., Anshary, A., Cyio, B., & Wanger, T. C. (2021). Landscape and farm-level management for conservation of potential pollinators in Indonesian cocoa agroforests. Biological Conservation, 257, 109106. 10.1016/J.BIOCON.2021.109106

Tscharntke, T., Clough, Y., Bhagwat, S. A., Buchori, D., Faust, H., Hertel, D., Hölscher, D., Juhrbandt, J., Kessler, M., Perfecto, I., Scherber, C., Schroth, G., Veldkamp, E., & Wanger, T. C. (2011a). Multifunctional shade-tree management in tropical agroforestry landscapes - A review. Journal of Applied Ecology, 48(3), 619–629. 10.1111/j.1365-2664.2010.01939.x

Tscharntke, T., Clough, Y., Bhagwat, S. A., Buchori, D., Faust, H., Hertel, D., Hölscher, D., Juhrbandt, J., Kessler, M., Perfecto, I., Scherber, C., Schroth, G., Veldkamp, E., & Wanger, T. C. (2011b). Multifunctional shade-tree management in tropical agroforestry landscapes - A review. Journal of Applied Ecology, 48(3), 619–629. 10.1111/j.1365-2664.2010.01939.x

Tscharntke, T., Clough, Y., Wanger, T. C., Jackson, L., Motzke, I., Perfecto, I., Vandermeer, J., & Whitbread, A. (2012). Global food security, biodiversity conservation and the future of agricultural intensification. Biological Conservation, 151(1), 53–59. 10.1016/J.BIOCON.2012.01.068

Waltert, M., Bobo, K. S., Sainge, N. M., Fermon, H., & Michael, M. (2005). From Forest to Farmland : Habitat Effects on Afrotropical Forest Bird Diversity. Ecological Applications, 15(4), 1351–1366.

Yodzis, P. (1988). The Indeterminacy of Ecological Interactions as Perceived Through Perturbation Experiments. Ecology, 69(2), 508–515. 10.2307/1940449

