## Appendix for "Food webs can deliver win-win strategies for tropical agroforestry and biodiversity conservation"

### APPENDIX I: Simulation study

*Data generation*

Our simulated study system consisted of 30 sites with varying environmental conditions. In these sites we simulated communities of 18 species in two trophic levels: predators (6 species) and prey (12 species). The dynamics of these communities were driven by an interaction matrix (**A**) which was shared across sites, and species-specific growth rates (**b**) that varied according to spatial and temporal environmental covariates. The parameters contained within **b** and **A** determined the equilibrium state of each community, which could differ between sites according to their respective covariates.

We started by assigning equilibrium biomasses for each species corresponding to a baseline of environmental covariates. The values of these equilibrium biomasses were not central to this exercise, but nevertheless we based them on our biological intuition of the system. We assigned baseline intrinsic growth rates $\nu_{i0}$ by assuming positive growth rates for prey and negative for predators (Table S1). In the absence of all other species the exponential of negative growth rates would result in a per-capita rate of change of $\exp(L_{ti}) < 1$, whilst a positive growth rate would result in $\exp(L_{ti}) > 1$. We considered intrinsic growth rates as a function of two covariates: one site-level covariate and one temporally varying covariate

(Table S1), so that intrinsic growth rate of species *i* at each site *f* and timepoint *t* was modelled as:

$$b_{ift} = \nu_{i0} + \nu_{i1}X_{1f} + \nu_{i2}X_{2t} \qquad \text{Equation S1}$$

We modelled regression coefficients as normally distributed (Table S1).

We then parametrised the **A** matrix as follows: first, all interspecific interaction terms other than those relating to predator prey relationships were assumed to be 0; second, the terms describing the effect of prey on predator were generated assuming a Holling type I functional response, where $a_{ij}$ was modelled as gamma distributed (Table S1). To reflect natural variation in trophic connectivity between species, we randomly set some interaction terms to 0, by multiplying each $a_{ij}$ term by a Bernoulli trial with probability $p = 0.6$, so that 29 out of 72 (12 prey x 6 predators) possible predator-prey interactions were 0. Third, the terms describing the effect of predator on prey, were calculated as $a_{ji} = -a_{ij}/\varepsilon_i$, where $\varepsilon_i$ was a trophic efficiency term (we set $\varepsilon_i = 0.5$; Table S1). Consequently, the effect of predator on prey was larger than the effect of prey on predator. Finally, we calculated the diagonals of the **A** matrix ($a_{ii}$ terms) by solving Equation 3 for the system equilibrium point (Table S1), so that

$$a_{ii} = -(\nu_{i0} + \textstyle\sum_{j \neq i} a_{ij} N_j^*)/N_i^* \qquad \text{Equation S2}$$

With this parametrisation, we generated data for biomass $N_{ti}$ over the 30 sites and 10 timepoints using Equations 1 - 3.

**Table S1.** Description of parameters used in simulation, including distribution or formula used to generate data and prior given to model. For gamma distributions, the parameters presented are shape and rate, for normal distributions they are mean and SD.

| **Parameter** | **Description** | **Simulated from** | **Prior** |
|---|---|---|---|

| | | | |
|---|---|---|---|
| $\nu_{prey0}$ | Intercept of linear predictor of intrinsic growth rate for prey groups | Γ(9,6) | Γ(44,22) |
| $\nu_{predator0}$ | Intercept of linear predictor of intrinsic growth rate for predator groups | - Γ(25,8) | - Γ(25,17) |
| $\nu_{i1}$ | Effect of site-level covariate on intrinsic growth rate | N(0,0.1) | N(0,0.5) |
| $\nu_{i2}$ | Effect of visit-level covariate on intrinsic growth rate | N(0,0.2) | N(0,0.5) |
| $a_{predator,prey}$ | Interaction coefficient for prey on predator | $-a_{prey,predator}\ \varepsilon$ | Γ(0.25,25) |
| $a_{prey,predator}$ | Interaction coefficient for predator on prey | - Γ(1,100) | $-a_{predator,prey}/\varepsilon$ |
| $a_{i,i}$ | Density dependence coefficient | Equation 6 | Γ(0.25,25) |
| ε | Energy efficiency of predators | 0.5 | Γ(100,200) |

*Model fitting*

We fit the model to these generated data by running 3 chains for 500,000 iterations (50,000 sample x 10 thinning rate), with an adaptation period of 2,000 iterations and a burn-in of 20,000. The model took 40 hrs to run using 3 cores for each of the parallel chains. We assessed model convergence by visually inspecting chains and with the Gelman-Rubin R-hat diagnostic, with convergence presumed when R-hat < 1.1. Once the model finished running, we sampled the parameters by taking 1,000 random draws from their posteriors. These parameter draws were used to calculate **N*** (Equation 4 main text), and we then compared estimated parameters and **N*** with true simulation values. We used the mean and 95% credible intervals (CIs) of these draws to assess model accuracy and precision.

*Results*

The model estimated all relevant parameters with high accuracy and precision (Fig. S1).

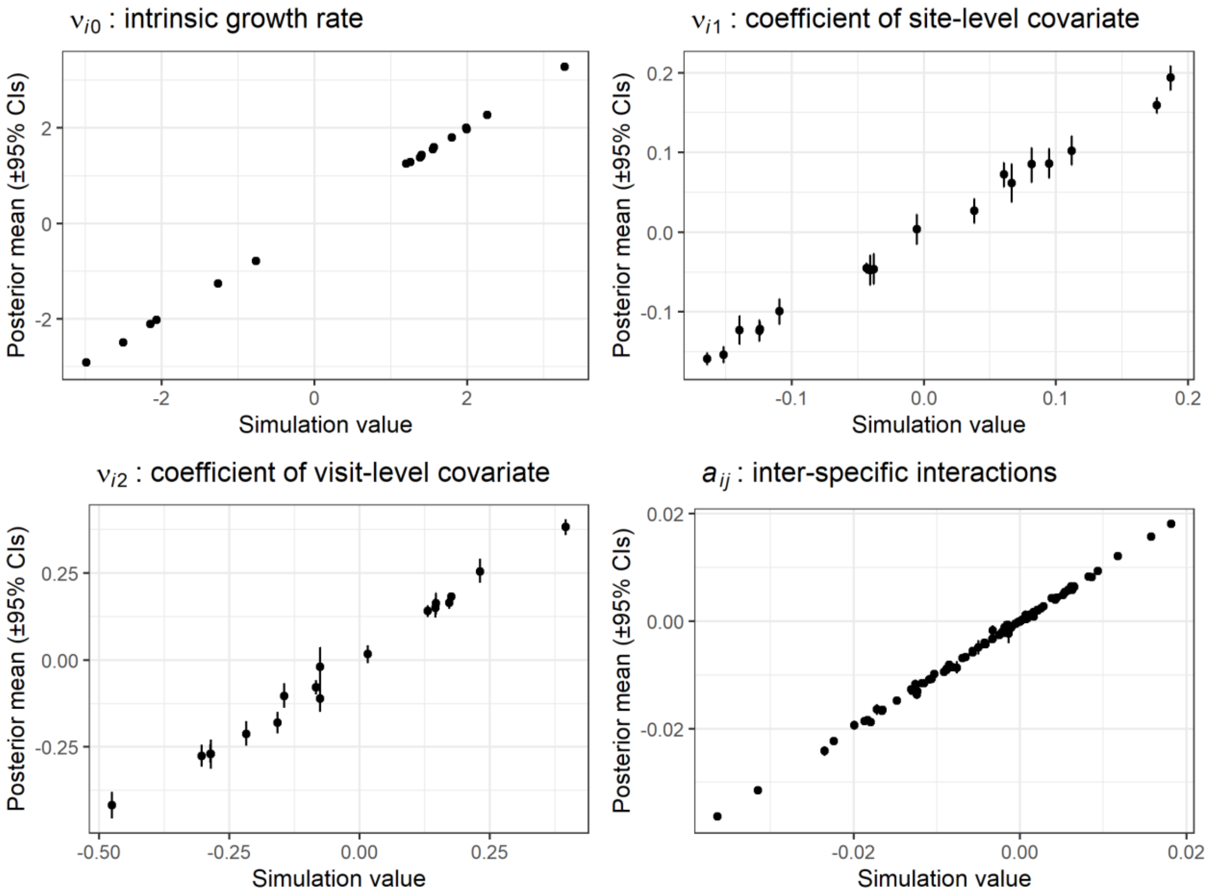


**Figure S1.** Correlation between parameter value used in simulation and parameter estimates from model. Parameter posteriors are represented using the median and 95% CIs of 1000 random draws from MCMC chains.

When we used parameter posteriors to estimate equilibrium biomasses of species, the retrieved biomasses were an accurate representation of the simulation biomasses (Fig. S2).

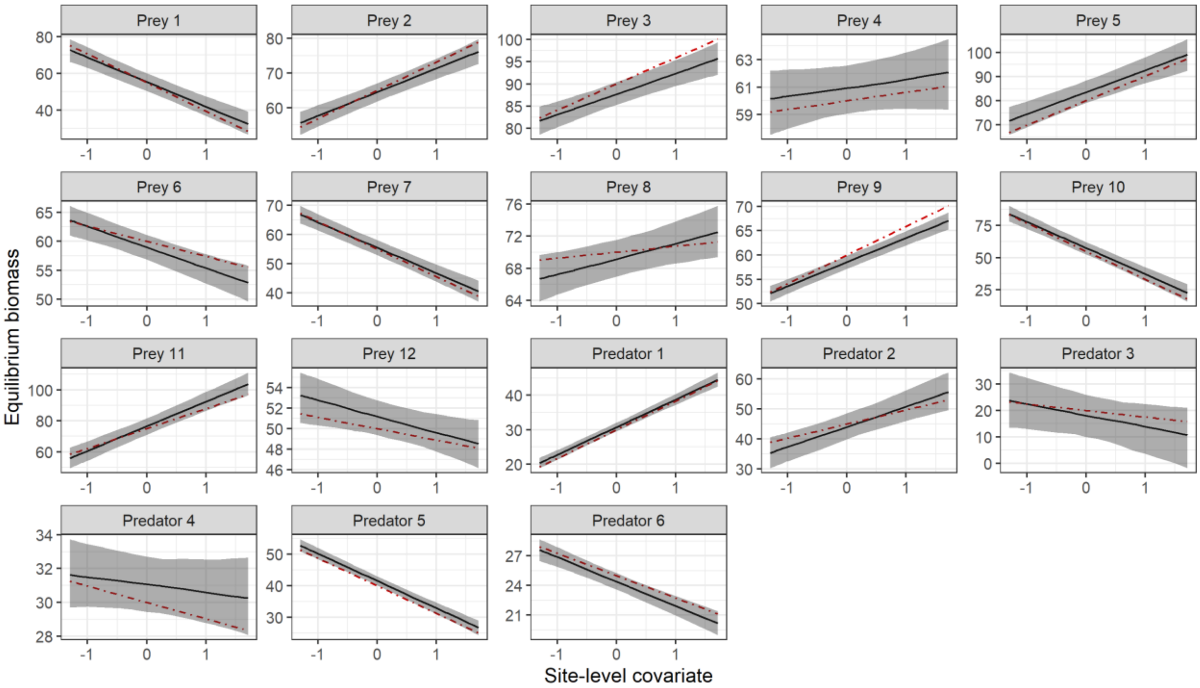


**Figure S2.** Effect of site-level covariate on true equilibrium biomasses (dashed lines) and equilibrium biomasses estimated from model predictions (solid lines and error shading). Equilibrium biomasses were calculated from 1000 random draws from parameter posteriors, and summarised using the mean and 95% CIs.

### APPENDIX II: Bird taxa in cocoa agroforestry

**Table S2.** Description of bird groups in community model, including species composition and brief information on ecology.

| **Group** | **Species** | **Description**[1] |
|---|---|---|
| Camaropteras | *Camaroptera brachyura, Camaroptera chloronota, Camaroptera superciliaris* | Small understory warblers that inhabits thickets in forest, forest edge and agricultural habitats. |
| Flycatchers | *Terpsiphone viridis, Terpsiphone rufiventer, Terpsiphone batesi,* | Medium-sized sallying flycatchers with long tails. Inhabit diverse |

| | | |
|---|---|---|
| | *Terpsiphone rufocinerea* | woodland and savanna habitats, as well as plantations and gardens. |
| Hylias | *Hylia prasina* | Stocky medium-sized warbler found in understory of forested habitats. |
| Kingfishers | *Ispidina picta, Ispidina lecontei* | Small kingfishers, mostly insectivorous. Not necessarily closely associated to water - also found in plantations and grasslands. |
| Wattle-eyes | *Platysteira castanea, Platysteira cyanea* | Small and plump flycatcher-like birds found in forested habitats. |
| Forest birds | *Alethe castanea, Bleda notatus, Criniger calurus, Criniger chloronotus, Criniger ndussumensis, Neocossyphus poensis, Phyllastrephus albigularis, Phyllastrephus icterinus, Phyllastrephus xavieri* | Large-bodied understory insectivores, generally restricted to primary or secondary forest. |

[1]Adapted from: del Hoyo, J., Elliott, A., Sargatal, J., Christie, D. A., & Kirwan, G. (2019). Handbook of the birds of the world alive. Lynx Edicions.

### APPENDIX III: Diet data from cocoa agroforestry

During field campaigns in Cameroon, we collected faecal samples from captured birds by placing individuals into a single-use paper bag that was in turn placed inside a regular cloth bird bag. Then, after the bird was processed, the paper bag was checked for faecal material. We collected any faecal material directly from the bag into a tube containing Longmire buffer

2% SDS. Samples were kept at room temperature throughout the sampling period, and then stored in the freezer at -20°C.

*Diet metabarcoding*

We extracted DNA from 442 faecal samples collected during the 2019-2020 fieldwork, of which 319 were successful. Alongside the faecal samples, we performed extractions on 21 control samples. We utilised between 0.06 and 0.08 g faecal matter (wet weight). DNA was extracted using a modified protocol of the EZNA Tissue Extraction kit (Omega Bio-tek). Samples were homogenised in a Tissueliser II (Qiagen) with 0.5g 0.5 mm silica-zirconia beads (BioSpec Products) and 650ul Gordon Buffer (0.1 M pH 8 Tris-HCl, 0.1 M pH 8 EDTA, 0.01 M NaCl and 0.01 M N-lauroylsarcosine) at 56°C per tube for 2 cycles of 5 mins at 25Hz, with samples rotated between cycles. Lysis continued for 12 to 24 hrs at 56°C in a water bath. Samples were then processed according to the manufacturer’s protocol, eluting in 70 uL Elution Buffer. Between three and five replicates of PCR were conducted for each sample to identify arthropod diet items. We used fwh2 primers (Vamos et al., 2017), which target an approximately 254 bp portion of the cytochrome oxidase I (COI) gene. Primers had a short section of the Illumina sequencing adapter appended to the end.

For each sample, the PCR replicated products were pooled in equal volumes and then cleaned using 0.7x 0.1% Sera-Mag carboxylate-modified SpeedBeads (ThermoScientific) following (Rohland & Reich, 2012) using 80% ethanol for washes. A second PCR was conducted using primers complementary to the overhang sequence and containing an individual specific pair of indices. Samples were then bead cleaned using 0.7x carboxyl-modified beads as above, quantified with the Qubit Broad Range dsDNA kit, pooled at 20nM, and sequenced on the Illumina MiSeq platform to produce 150 bp paired-end sequences.

Raw sequences processed following the methods described in (Jarrett et al., 2020). Briefly, the sequences were quality trimmed using Sickle v1.33 (Joshi & Fass, 2011), and error corrected following Schirmer et al. 2015 using BayesHammer (Nikolenko et al., 2013) through the SPAdes program v3.10.1 (Bankevich et al., 2012). Sequences were merged using PEAR v0.9.6. (Zhang et al., 2014) and PCR primers were trimmed off using CutAdapt v1.10 (Martin, 2011). Chimeric sequences were identified via usearch v11.0.667 (Edgar, 2010) implemented through a modified version of the program DAMe (Zepeda-Mendoza et al., 2016) and removed. DAME was used to prepare for OTU clustering, and to filter sequences ± 30bp of the target sequence length (without primers). Sequences were then clustered into molecular operational taxonomic units (OTUs) at the 97% identity level using Sumaclust (Mercier et al., 2013). Only OTUs with ≥ 5 sequences were retained. Following Alberdi et al. (2018) and Aizpurua et al. (2018), we assigned taxonomy via a BLAST search of the Genbank NT database. Using blastn in BLAST+ 2.7.1 we retrieved the best 20 matches for each representative OTU sequence. Using custom python and R scripts (Aizpurua et al., 2018; Alberdi et al., 2018), we retained the highest common taxonomic information among matches with the maximum bitscore, and assigned taxonomy to each OTU based on identity: For matches with ≥95% identity we assigned order- level taxonomy; ≥96.5% we assigned family-level, and for ≥98% we assigned genus and species-level taxonomy.

After taxonomic assignment, we performed several data cleaning steps: we excluded all OTUs that were detected at <1% of the total number of reads, we excluded all OTUs that appeared in the control samples at >1% of the total number of reads, and finally, we considered only OTUs of the class Insecta and Arachnida (to focus on diet items rather than, e.g., fungal species).

*Data analysis*

The objective of this analysis was to identify prey groups rarely consumed by predator taxa, in order to thin out interaction parameters in the community model. Using the diet metabarcoding dataset resulting from the steps above, we calculated for each individual bird the proportion of reads made up by each arthropod group in the food web model, by clustering OTUs across groups. We then grouped birds according to the food web clusters (camaroptera, flycatcher, wattle-eye, hylia, kingfisher, forest birds) and calculated the mean proportion across individuals (Fig. S3). We considered that if a certain diet item constituted on average <5% of the reads of a given bird group, that trophic parameter was set to 0 in the community model (Fig. S3). We made several important adaptations to this rule: first, several of the arthropod orders in the community model occurred as two separate nodes, one representing pests and the other non-pests. In the diet data, we did not have this distinction, as many OTUs were not identified to species level and therefore would not allow us to distinguish whether they corresponded to pests or not. Therefore, we used the same primers for interactions between a given bird group and the pest and non-pest factions; for instance, if we detected <5% reads of Coleoptera in the diet of a certain bird group, we set to 0 the parameters relating to the interaction between this bird group and both Coleoptera pests and non-pests.

Second, likely due to the chosen primer, Araneae OTUs occurred at low numbers of reads in our dataset across all individuals. Following the analysis described above, no bird taxa contained Araneae at >5% mean proportion of reads (Fig. S3). Nevertheless, we know from our arthropod survey data and previous studies (Ferreira et al., 2023; Jarrett et al., 2023) that Araneae are not uncommon in our system, and likely consumed by insectivorous birds (Garfinkel et al., 2022; Jarrett et al., 2020). Therefore, for Araneae we did not set to 0 the trophic parameters corresponding to the bird groups that consumed them at highest frequency, which were wattle-eyes (4.1%) and kingfishers (3.7%; Fig. S3).

Finally, a similar situation occurred with brown capsid: as this was the only species-level prey node in the community model, it was likely to occur in the diet at much lower proportions than a node that includes a whole insect order. We therefore considered the 5% cut-off inappropriate for brown capsid. Instead, we considered that a trophic link existed between bird taxa and brown capsid for any bird group which, according to the diet metabarcoding data, consumed brown capsid at no matter what frequency. This was true for kingfishers, flycatchers and forest birds.

Overall, this analysis resulted in the removal (parameter set to 0) of 25 trophic connections between birds and arthropods.

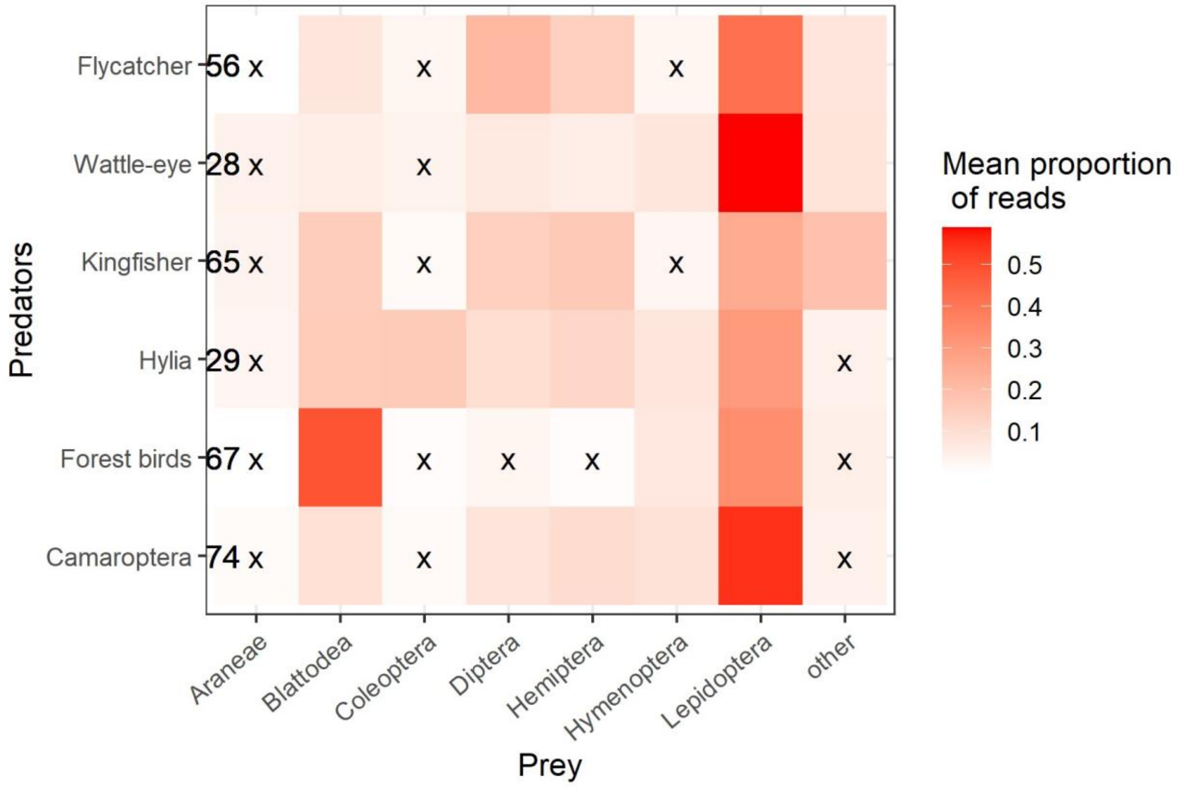


**Figure S3.** From diet metabarcoding data, the mean proportion of reads of each prey group in the diet of each predator group. Sample sizes are indicated next to the predator group names. White squares with an ‘x’ indicate an average of <5% proportion of reads of this prey group in the samples.

### APPENDIX IV: Observation models

Birds: we modelled bird population size by integrating mist-net captures and acoustic recordings into a joint likelihood, as in Jarrett et al. (2022). This model assumes that there was an underlying population of birds at each site, which was sampled by both methods. The only modification we made to the Jarrett et al. (2022) model was the grouping of species, as we were interested in a specific set of taxa (Fig. 2 main text). For the observation model, we used the whole dataset described in Jarrett et al. (2022), to maximise sample sizes for shared parameter estimation, but then distinguished the subset of groups targeted in the process model. We assumed an equal mist-net capture rate for camaroptera, hylia, kingfishers, wattle-eyes and flycatchers as they are all small, morphologically similar genera. We allowed for distinct vocalisation rates between these genera.

Arthropods: we modelled arthropod population sizes by integrating data from three common survey techniques, as in Jarrett et al. (2023).

We ran both observation models with 3 chains for 20,000 MCMC iterations (plus 10,000 burn-in). Both observation models estimated the population size of taxa, so we converted population size to biomass by correcting for body mass and effective area sampled. For birds, we calculated average body mass for each taxon from field data. For arthropods, we assumed a mean body mass of 0.01 g across taxa (Byrne et al., 1988; Hancock & Legg, 2012). Mean biomass and standard deviation were then provided to the community model as data (Fig. S4).

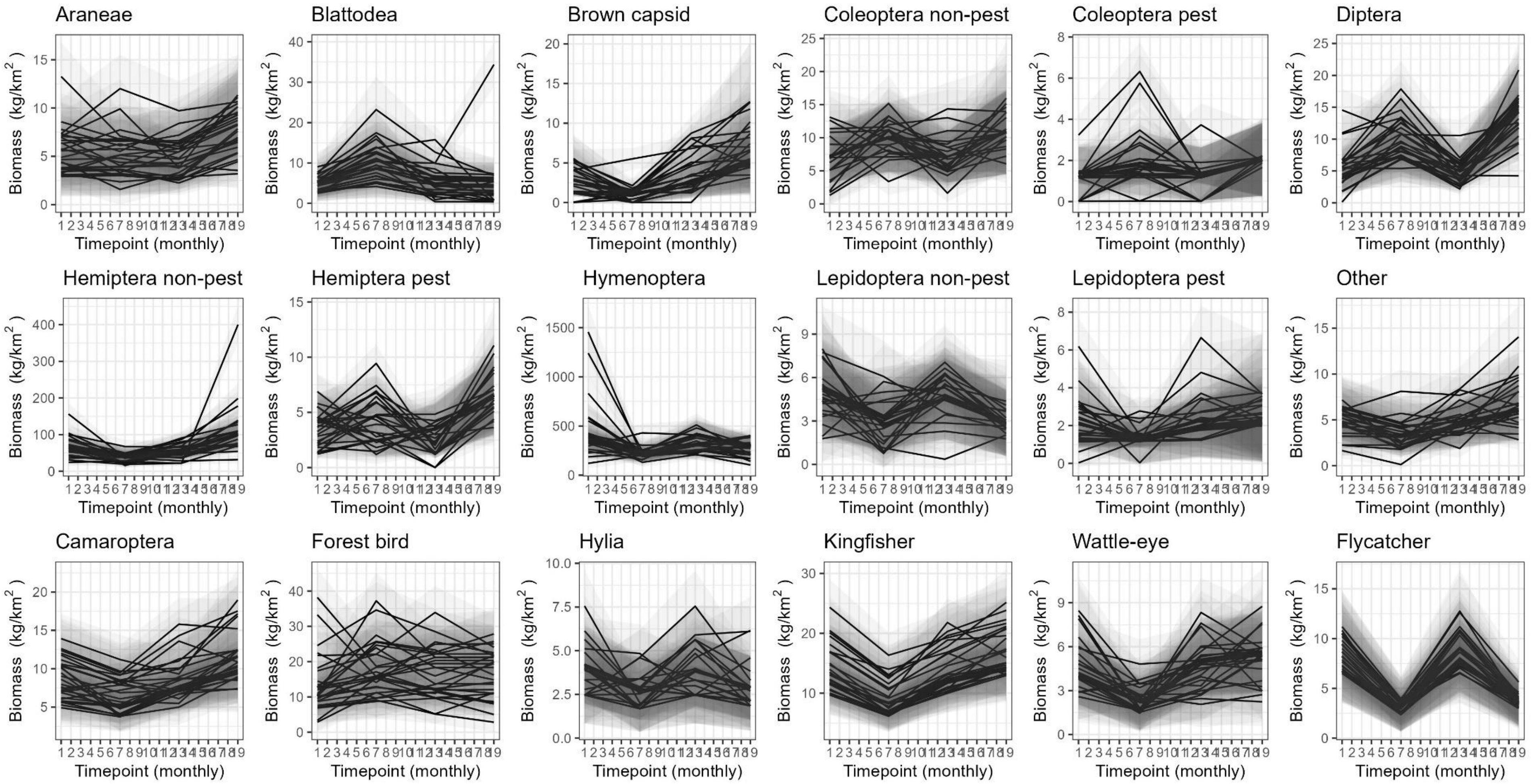


**Figure S4.** Biomass of each taxon in the community model through time, as estimated by observation models. Lines represent mean posterior distribution, and shaded areas are standard deviation of posteriors. These data were provided to the process model.

### APPENDIX V: Additional results from cocoa agroforestry

*Parameter estimates*

Parameters relating to groups' growth rates and interactions are presented in Figure S5. Intrinsic growth rates of arthropod groups ranged from 1.18 in Coleopteran pests to 0.16 in Hymenopterans, which is equivalent to a population doubling time in the range of 17 days to 4 months. The intrinsic growth rates of birds ranged from -0.19 for forest birds to -0.38 for hylia, which is equivalent to a population half-life of 109 to 54 days.

The effect of season and shade cover on growth rates varied between taxa (Fig. S5). Araneae, Blattodea, Diptera, Hymenoptera and forest birds showed a positive trend between growth rate and shade cover, whilst brown capsid, Hemiptera non-pest, Lepidoptera pest, camaroptera, kingfisher and flycatchers showed a negative trend. For season, Hemiptera non-pest, Hymenoptera, camaroptera, kingfisher, wattle-eyes and flycatcher had higher growth rates in the wet season, whilst Araneae, Blattodea, Coleoptera non-pest, Diptera, Hemiptera pest and Lepidoptera had higher growth rates in the dry season.

Interaction parameters ranged from -0.32 to 0.03, with the lowest values associated to density dependence parameters in certain taxa, as well as certain interactions between bird and arthropod taxa (Fig. 4 main text; Fig. S6). The $a_{ij}$ parameters represent consumption scaled by density; for consumption per se see Figure S7.

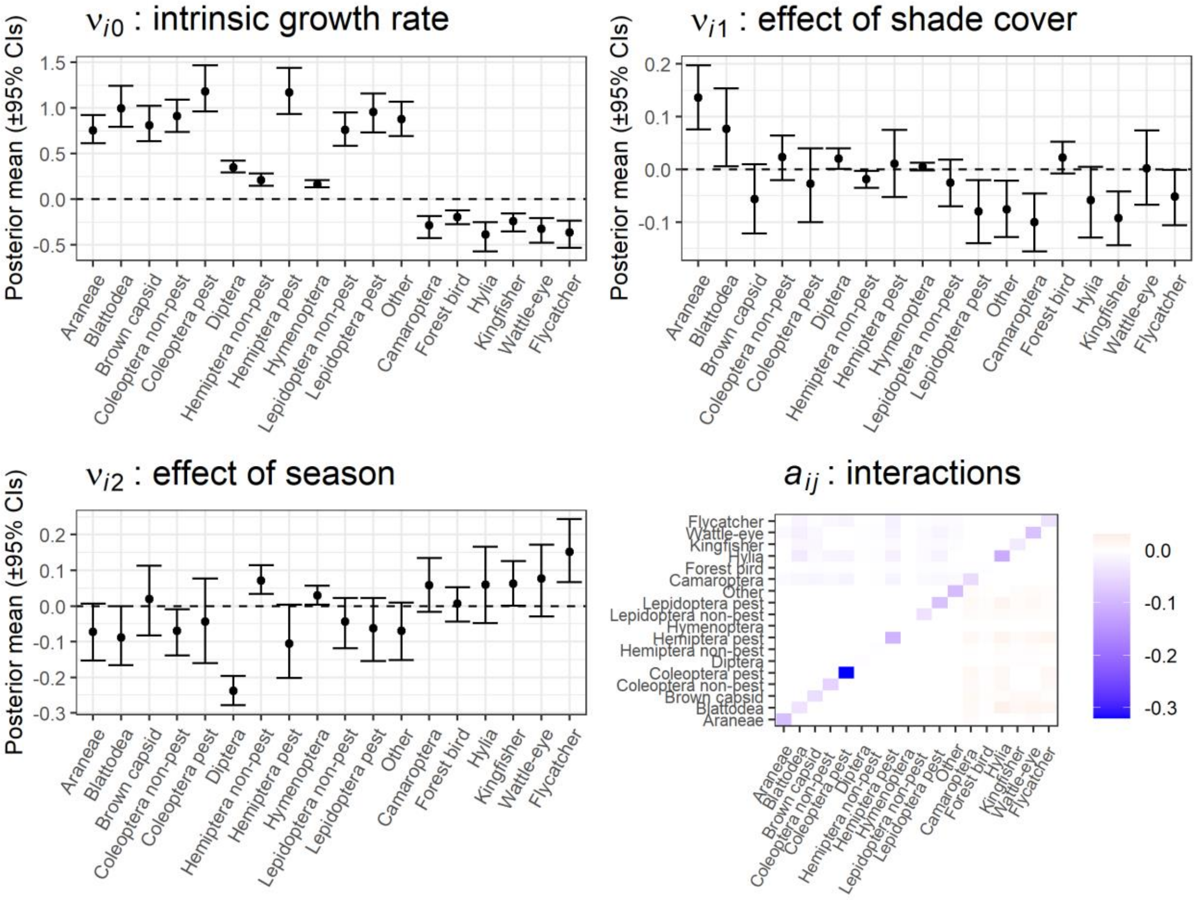


**Figure S5.** Growth rate and interaction parameters estimated by model. Parameter posteriors were summarised using the mean and 95% CIs of 1000 random draws from MCMC chains.

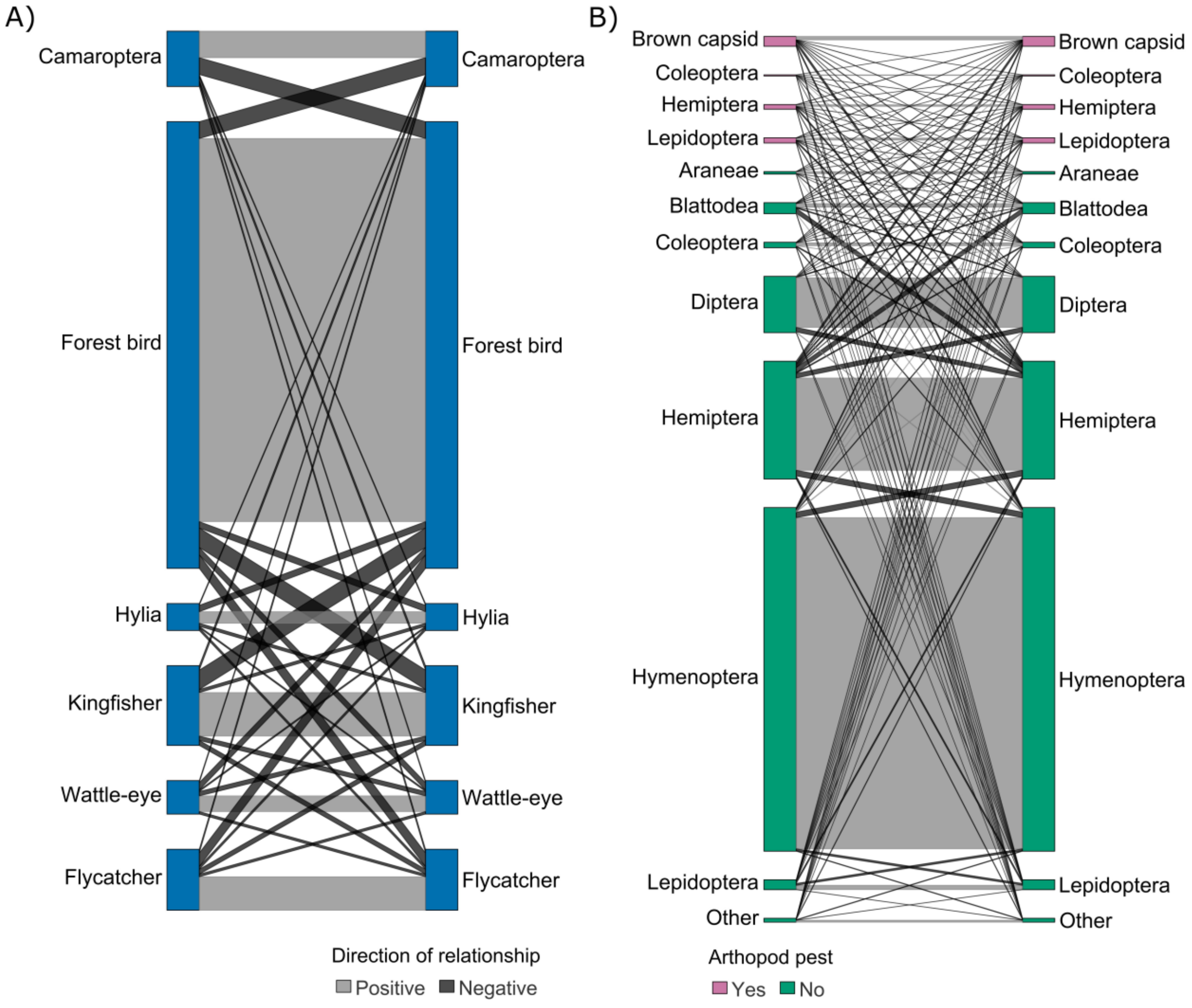


**Figure S6.** Interactions among bird taxa (A) and among arthropod taxa (B). Represented is the net effect of bird groups on arthropod groups, i.e., the change in equilibrium biomass of one taxon caused by the addition of biomass of another taxon.

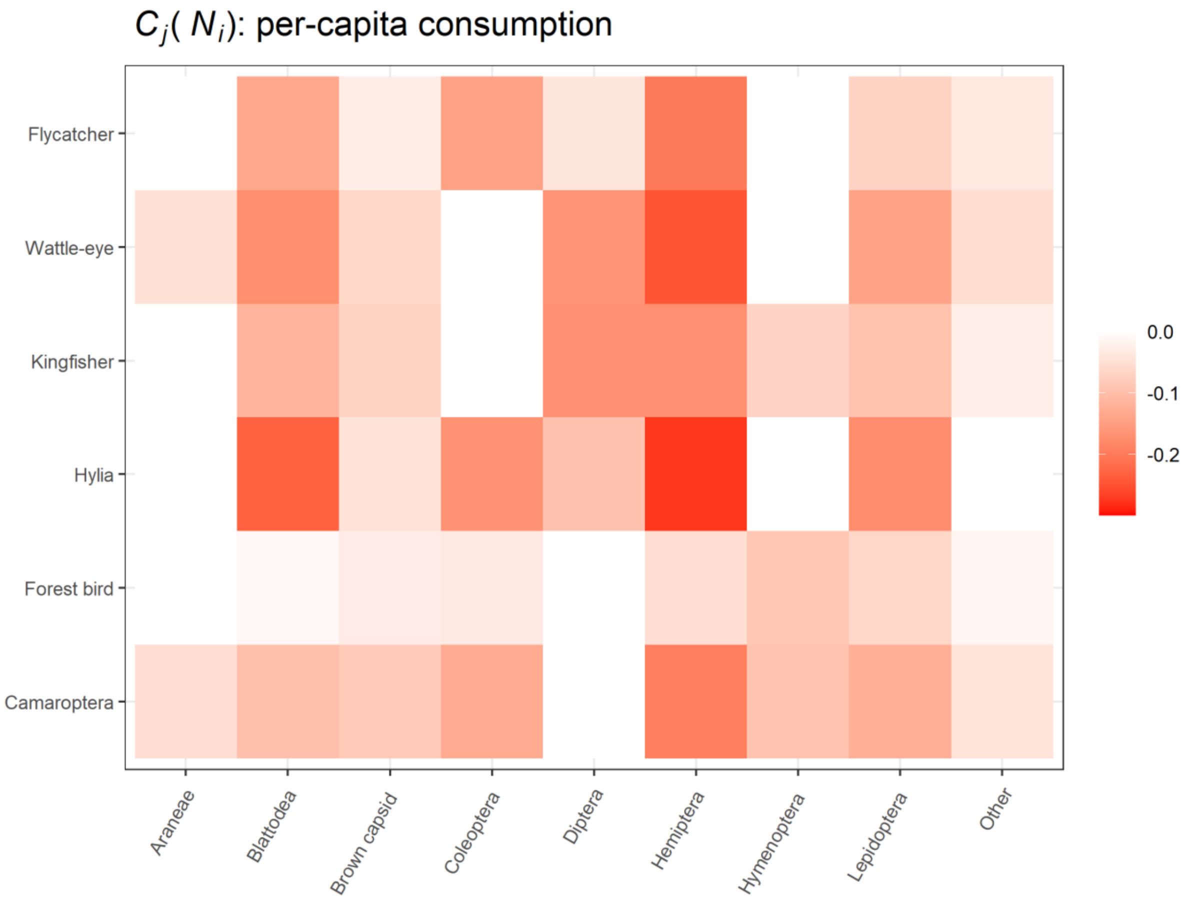


**Figure S7.** Per capita consumption of arthropod taxa by bird taxa, estimated from community model.

*Estimating equilibrium states*

We estimated equilibrium biomass analytically for each taxon at each site and season using Equation 4 (Fig. S6); equilibrium biomasses match the composition at which communities settle when data are simulated stochastically from parameters.

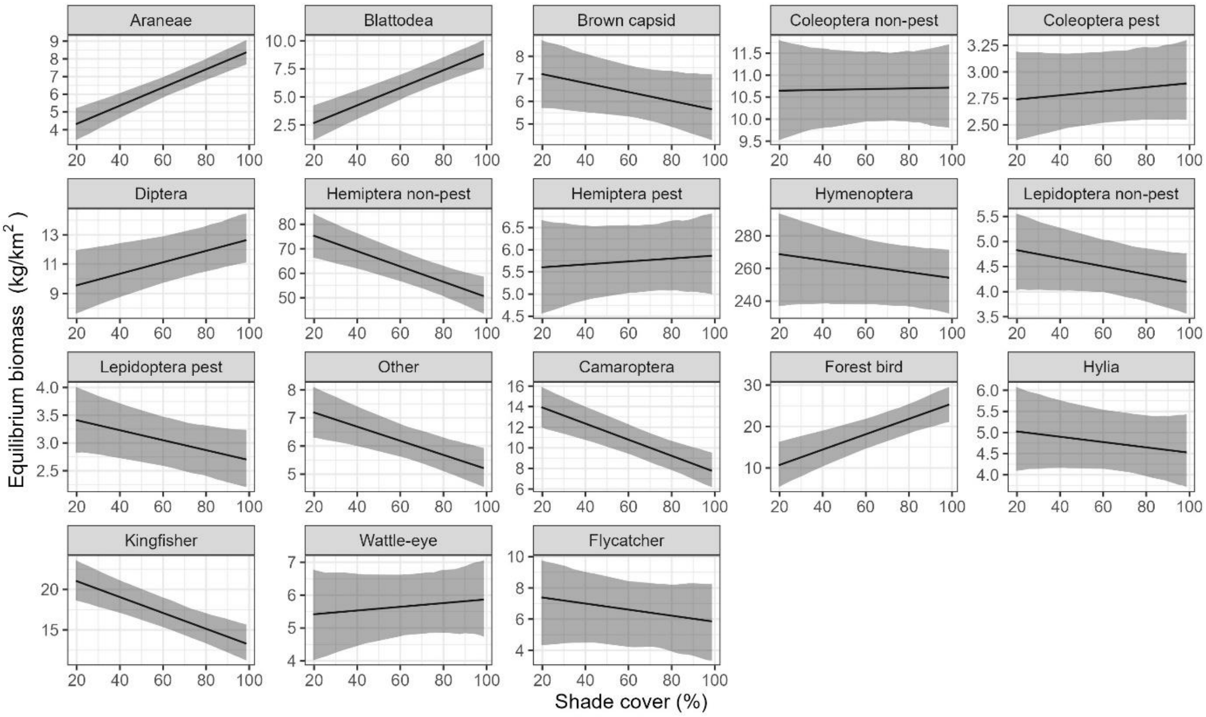


**Figure S8.** Effect of shade cover and season on equilibrium biomass of each group estimated from model parameters. Presented here is the equilibrium biomass corresponding to the dry season – the model estimates also rainy season equilibria.

### REFERENCES

Aizpurua, O., Budinski, I., Georgiakakis, P., Gopalakrishnan, S., Ibañez, C., Mata, V., Rebelo, H., Russo, D., Szodoray-Parádi, F., Zhelyazkova, V., Zrncic, V., Gilbert, M. T. P., & Alberdi, A. (2018). Agriculture shapes the trophic niche of a bat preying on multiple pest arthropods across Europe: Evidence from DNA metabarcoding. *Molecular Ecology*, *27*(3), 815–825. https://doi.org/10.1111/mec.14474

Alberdi, A., Aizpurua, O., Gilbert, M. T. P., & Bohmann, K. (2018). Scrutinizing key steps for reliable metabarcoding of environmental samples. *Methods in Ecology and Evolution*, *9*(1), 134–147. https://doi.org/10.1111/2041-210X.12849

Bankevich, A., Nurk, S., Antipov, D., Gurevich, A. A., Dvorkin, M., Kulikov, A. S., Lesin, V. M., Nikolenko, S. I., Pham, S., Prjibelski, A. D., Pyshkin, A. V, Sirotkin, A. V, Vyahhi, N., Tesler, G., Alekseyev, M. A., & Pevzner, P. A. (2012). SPAdes: a new genome assembly algorithm and its applications to single-cell sequencing. *Journal of Computational Biology : A Journal of Computational Molecular Cell Biology*, *19*(5), 455–477. https://doi.org/10.1089/cmb.2012.0021

Byrne, D. N., Buchmann, S. L., & Spangler, H. G. (1988). Relationship Between Wing Loading, Wingbeat Frequency and Body Mass in Homopterous Insects. *Journal of Experimental Biology*, *135*(1), 9–23. https://doi.org/10.1242/JEB.135.1.9

Edgar, R. (2010). *Usearch*. Lawrence Berkeley National Lab.(LBNL), Berkeley, CA (United States).

Ferreira, D. F., Jarrett, C., Wandji, A. C., Atagana, P. J., Rebelo, H., Maas, B., & Powell, L. L. (2023). Birds and Bats Enhance Yields in Afrotropical Cacao Agroforests Only Under High Shade. *Agriculture, Ecosystems & Environment*, *345*, 108325.

Garfinkel, M., Minor, E., & Whelan, C. J. (2022). Using faecal metabarcoding to examine consumption of crop pests and beneficial arthropods in communities of generalist avian insectivores. *Ibis*, *164*, 27–43. https://doi.org/10.1111/ibi.12994

Hancock, M. H., & Legg, C. J. (2012). Pitfall trapping bias and arthropod body mass. *Insect Conservation and Diversity*, *5*(4), 312–318. https://doi.org/10.1111/J.1752-4598.2011.00162.X

Jarrett, C., Cyril, K., Haydon, D., Wandji, A., Ferreira, D., Welch, A., Powell, L., & Matthiopoulos, J. (2023). Fewer pests and more ecosystem service-providing arthropods in shady African cocoa farms: Insights from a data integration study. *Journal of Applied Ecology, In press*.

Jarrett, C., Haydon, D. T., Morales, J. M., Ferreira, D. F., Forzi, F. A., Welch, A. J., Powell, L. L., & Matthiopoulos, J. (2022). Integration of mark-recapture and acoustic detections for unbiased population estimation in animal communities. *Ecology*, *103*(10), e3769.

Jarrett, C., Powell, L. L., McDevitt, H., Helm, B., & Welch, A. J. (2020). Bitter fruits of hard labour: diet metabarcoding and telemetry reveal that urban songbirds travel further for lower-quality food. *Oecologia*, *193*(2), 377–388. https://doi.org/10.1007/s00442-020-04678-w

Joshi, N. A., & Fass, J. (2011). *Sickle: A sliding-window, adaptive, quality-based trimming tool for FastQ files (Version 1.33)[Software]*.

Martin, M. (2011). Cutadapt removes adapter sequences from high-throughput sequencing reads. *EMBnet. Journal*, *17*(1), 10–12.

Mercier, C., Boyer, F., Bonin, A., & Coissac, E. (2013). SUMATRA and SUMACLUST: fast and exact comparison and clustering of sequences. *Programs and Abstracts of the SeqBio 2013 Workshop*, 27–29.

Nikolenko, S. I., Korobeynikov, A. I., & Alekseyev, M. A. (2013). BayesHammer: Bayesian clustering for error correction in single-cell sequencing. *BMC Genomics*, *14*(1), 1–11. https://doi.org/10.1186/1471-2164-14-S1-S7/TABLES/3

Rohland, N., & Reich, D. (2012). Cost-effective, high-throughput DNA sequencing libraries for multiplexed target capture. *Genome Research*, *22*(5), 939–946. https://doi.org/10.1101/gr.128124.111

Vamos, E. E., Elbrecht, V., & Leese, F. (2017). Short COI markers for freshwater macroinvertebrate metabarcoding. *Metabarcoding and Metagenomics 1: E14625*, *1*, e14625-. https://doi.org/10.3897/MBMG.1.14625

Zepeda-Mendoza, M. L., Bohmann, K., Carmona Baez, A., & Gilbert, M. T. P. (2016). DAMe: a toolkit for the initial processing of datasets with PCR replicates of double-tagged amplicons for DNA metabarcoding analyses. *BMC Research Notes*, *9*(1), 1–13.

Zhang, J., Kobert, K., Flouri, T., & Stamatakis, A. (2014). PEAR: a fast and accurate Illumina Paired-End reAd mergeR. *Bioinformatics*, *30*(5), 614–620. https://doi.org/10.1093/bioinformatics/btt593
